# A versatile and upgraded version of the LundTax classification algorithm applied to independent cohorts

**DOI:** 10.1101/2023.12.15.571519

**Authors:** Elena Aramendía Cotillas, Carina Bernardo, Srinivas Veerla, Fredrik Liedberg, Gottfrid Sjodahl, Pontus Eriksson

## Abstract

Stratification of cancer into biologically and molecularly similar subgroups is a cornerstone of precision medicine and transcriptomic profiling has revealed that urothelial carcinoma (UC) is a heterogeneous disease with several distinct molecular subtypes. The Lund Taxonomy classification system for urothelial carcinoma aims to be applicable across the whole disease spectrum including both non-muscle invasive and invasive bladder cancer. For a classification system to be useful it is of critical importance that it can be applied robustly and reproducibly to new samples. Many transcriptomic methods used for subtype classification are affected by the choice of expression platform, data preprocessing, cohort composition, and tumor purity. The application of a subtype classification system across studies therefore comes with a degree of uncertainty regarding whether the predictions in a new cohort accurately recapitulate the originally intended stratification. Currently, only limited data has been published evaluating the transferability and applicability of existing stratification systems and their respective classification-algorithms to external datasets. In the present investigation we develop a single sample classifier based on in-house microarray and RNA-sequencing data intended to be broadly applicable across datasets, studies, and tumor stages. We evaluate the performance of the proposed method and the Lund Taxonomical classification across 10 published bladder cancer cohorts (n=2560 cases) by examining the expression of characteristic subtype associated gene signatures, and whether complementary data such as mutations, clinical outcomes, response, or variant histologies are captured by our classification. Effects of varying sample purity on the classification results were also evaluated by generating low-purity versions of samples in silico. We show that the classifier is robustly applicable across different gene expression profiling platforms and preprocessing methods, and less sensitive to variations in sample purity.

The classifier is available as the ‘LundTaxonomy2023Classifier’ R package on [GitHub](https://github.com/LundBladderCancerGroup/LundTaxonomy2023Classifier).

## Introduction

Molecular subtyping aims to classify and categorize similar tumor samples within a broader group based on their distinct molecular characteristics. Subtype stratifications help us gain a better biological understanding of the disease and may enable more precise clinical decision making. While stratification schemes developed in isolated works of research can provide new insights and research-hypotheses, being able to extend a classification strategy to new datasets is essential to validate, expand, and strengthen the results. However, there are many technical and biological factors that make this a challenge. Transcriptomic research cohorts have been produced using a number of microarrays or RNA-sequencing methods, each using different labeling kits, library preparation techniques, sequencing technology, and data preprocessing methods. Studies also differ in biopsy sample origin e.g., cystectomy or transurethral resection of bladder tumors (TURB) specimens, and tissue preservation e.g., fresh frozen tissue or paraffin embedded tissue. Furthermore, cohort design is usually driven by particular clinical research questions, resulting in cohorts enriched for a selected subset of tumors. Consequently, it may be challenging to generalize results to a broader disease spectrum, especially when results are derived from relative gene expression values where the data has been row-centered or scaled across the cohort and therefore strongly dependent on the composition of the cohort. If a classifier is trained on such relative data, its application is only appropriate to new data that has been preprocessed and row-centered to reproduce the relative differences of the training-cohort. This can be difficult to achieve if a new cohort has a significantly different composition, or if the new data is different from that of the original cohort (e.g., microarray vs RNA-sequencing). To overcome this issue, recent studies have designed single-sample predictors (SSP) intended to be applied to individual samples by using some form of raw non-cohort normalized data (1–6). While this approach circumvents issues related to cohort composition, to our knowledge there has been no systematic validation of such prediction methods across external data. Moreover, molecular subtyping is dependent on the principles used to define cancer subtypes, and there is currently no strict definition of what constitutes a subtype. In general terms, a subtype denotes a group of objects that share common features, exhibit similarities, and differ from other groups of objects. Subtype assignments can be based on specific molecular characteristics, such as the gene fusion BCR/ABL in chronic myeloid leukemia. Alternatively, they could be based on complex traits, such as altered transcriptional programs as measured by microarrays or RNA-sequencing. Given the numerous features that can be used to categorize a tumor, it is essential to carefully consider the similarities and differences captured by particular classification systems (7). It is also important to consider whether the goal is to group cancers based on the properties of the entire biopsy or the properties specific to the cancer cells proper. A common approach has been to perform hierarchical clustering of gene expression data obtained from bulk biopsies to define subgroups. We have, however, repeatedly observed that the derived cluster structures can be largely shaped by biopsy purity. The content of immune and stromal cells, as well as the level of proliferation within the biopsy results in strong coherent expression signatures that often overshadow and mask differences within the cancer cells proper (8). Through extensive gene expression cluster analyses paired with immunohistochemical (IHC) analyses using antibodies for 25 proteins, we could evaluate and refine the Lund Taxonomy and reveal discrepancies between the actual cancer cell phenotypes as identified by IHC, and groupings based on global mRNA clustering (8). We then adopted a supervised approach and trained the RNA-based algorithm to identify IHC determined cancer cell phenotypes. This allowed us to resolve gene expression groupings caused by varying levels of infiltration and proliferation and showed that cancer cell phenotypes may be detected using RNA-based analyses only (9). In the present work we have refined and made publicly available a Random Forest based single-sample gene expression predictor (10) that classifies bladder cancer samples into the five main Lund Taxonomy cancer cell subtypes Urothelial-like (Uro), Genomically unstable (GU), Basal squamous-like (Ba/Sq), Mesenchymal-like (Mes-like), and Small cell Neuroendocrine-like (Sc/NE), and subclassifies Uro further into UroA, UroB, and UroC (Figure 1A) (8, 9, 11, 12). We have applied this classification algorithm to several independent and published datasets of both muscle invasive and non-muscle invasive urothelial carcinomas. The algorithm was robustly applicable across different gene expression profiling platforms and preprocessing methods, and less sensitive to variations in sample purity.

**Fig. 1.**
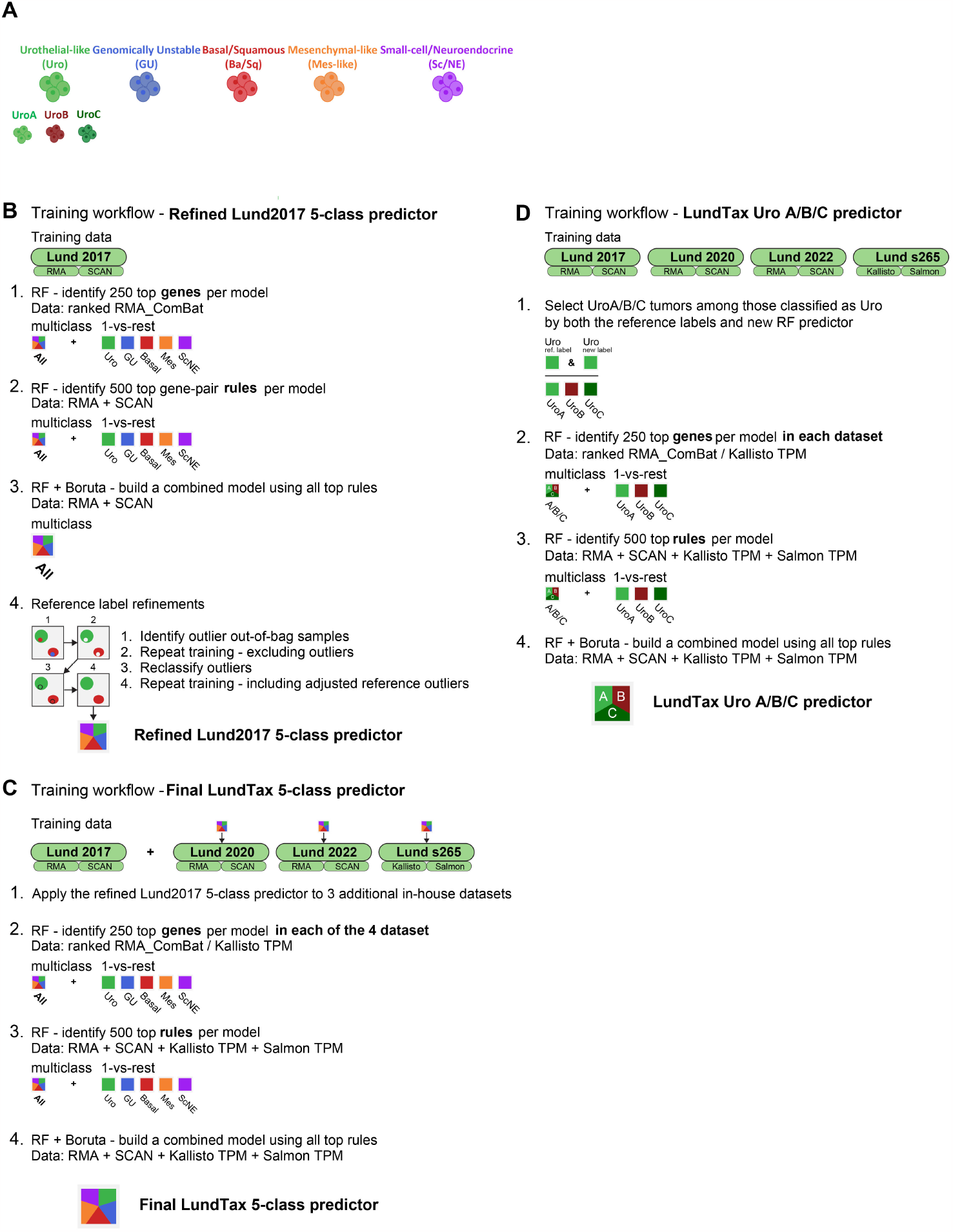
**A:** Schematic representation of the five major LundTax molecular subtypes and the subdivision of the urothelial-like into UroA, UroB, and UroC. The respective color codes are used all throughout the manuscript. **B-D:** Flowchart showing the major steps in the development of the LundTax2023 algorithm, including a first a Lund2017-based 5-class classifier (**B**), the full 5-class classifier (**C**) and the Uro A/B/C predictor (**D**).

## Methods

### Microarray data preprocessing

Affymetrix microarray datasets were normalized with two preprocessing pipelines: RMA (Robust Multichip Average) and SCAN (Single-channel array normalization) using the R packages oligo v3.17 (13) and SCAN.UPC v3.18 (14), respectively. RMA is a widely used cohort-based Affymetrix array preprocessing method, while SCAN is a single sample normalization approach designed to counter the impact of cohort composition on the normalization process and is used by a commercial provider of Affymetrix-based bladder cancer transcriptomic profiling. We used BrainArray V25 probe annotations based on Gencode 36 across all Affymetrix microarrays (Exon, Gene) (15), providing a static set of annotation files that also avoids multi-mapping probes, giving a readily reproducible gene expression summarization directly at the gene level. Additionally, “gene_ biotype”, “hgnc_ symbol”, and “hgnc_ id” was added to each array annotation using the biomaRt R package (16) using Ensembl 108/Gencode42. A small number of features (less than five on either array) were discarded due to a non-unique HGNC gene symbol identifier.

The final array datasets were filtered to retain protein coding genes with HGNC gene names.

### RNA-sequencing data preprocessing

RNA-sequencing datasets with raw FASTQ files available were processed using the pseudoaligners Kallisto (v0.48.0) (17) and Salmon (v1.9.0) (18) with GRCh38 Gencode39 reference indexes. To improve the stability of the TPM data for visualization, we adopted a transcript filtering step for both the Kallisto and Salmon output files, removing technically variable transcripts (including microRNA, miscellaneous-RNA, TCR and BCR coding genes, V, D, and J regions, histone-coding and mitochondrial genes) (19). Retained transcripts were then rescaled and summarized to gene level in TPM format, as well as GeTMM format (20) using the R package txImport v1.26.1 (21) and a fixed tx2gene file. All RNA-seq datasets with available raw data were additionally quantified with Kallisto using a Gencode23 reference from the Toil pipeline (22). For external datasets with only preprocessed data available, we applied the classification algorithm directly on the available published TPM or FPKM values. If any genes were missing from these datasets, this was addressed by updating outdated gene symbol annotation and through the k-Nearest Neighbors imputation feature of multiclassPairs (23). As the prediction method is applied to individual samples, TPM and FPKM can be used interchangeably as these have a perfect Spearman correlation of 1, given that they are calculated from the same counts and gene lengths for a sample.

### In-house datasets

In total, four cohorts were used to train the new model. Three datasets were generated on the Affymetrix Human Gene 1.0 ST microarray platform and largely correspond to previously published cohort datasets. The Lund2017 (8) dataset included 301/307 samples from GSE83586, excluding 6 samples labeled as “infiltrated”, the Lund2020 (24) dataset contained 173 samples from GSE128959, and the Lund2022 (25, 26) series contained 310 samples in total, 117 included in GSE169455, 37 in GSE222073 and 156 unpublished samples. All microarray data was reprocessed from the original CEL files. The s265 cohort contained 265 RNA-sequenced tumors of mixed stages. Libraries were prepared using TruSeq Stranded library preparation kit and were sequenced on the Illumina NextSeq 500 system (75bp paired-end). The data was quantified using Kallisto and Salmon. The full set of CEL files, Kallisto and Salmon output files, and processed data (as used in the study) has been deposited (10.5281/zenodo.10362517). Preprocessing code and transcript to gene mappings have also been deposited. Sample IDs, reference labels, intermediate training labels, IHC labels, and final subtype labels for all utilized training samples are listed in Supplementary Table 1. Based on previous characterizations, the transcriptomic subtype label could also receive the suffix “-inf” (e.g., “GU-Inf”) to denote samples with higher infiltration. This distinction was not utilized in the current study, except for the sub stratification of Uro samples where Uro-Inf classified samples were excluded from training as they did not have an assigned UroA/B/C label.

### Immunohistochemistry

The three array datasets had complementary tissue microarray IHC, performed as described in the respective previous publications (8, 24, 25). Briefly, two 1.0 mm cores from tumor-rich areas of the TUR-BT tissue blocks were embedded into tissue microarrays. TMA blocks were then sectioned (4 μm) and stained with various subtype informative IHC-markers. The minimal shared set of markers available for all three cohorts used here were; CDH1, EPCAM, GATA3, RB1, FGFR3, CCND1, KRT5, KRT14, TUBB2B, VIM, and ZEB2. Primary antibodies, incubation, and staining conditions were as described in the original studies. Tissue sections were stained using an Autostainer Plus (Dako, Glostrup, Denmark), scanned (AxioScan Z1, Zeiss, Oberkochen, Germany), and evaluated as digital images (Xplore v5.1, Philips, Amsterdam, Netherlands). Staining was evaluated either as intensity (0-3), as percentage (10 bins), or both (Intensity x Percentage), depending on the staining pattern and the mean marker score for the two cores was used, as described in the original studies.

### External datasets

External datasets were collected from public portals including the Gene Expression Omnibus (GEO), the European Genome Archive (EGA), Genomic Data Commons (GDC), and the Database of Genotype and Phenotype (dbGaP). The raw TCGA-BLCA RNA-sequencing dataset was obtained in FASTQ format and preprocessed using Kallisto and Salmon. Three independently preprocessed versions of the TCGA-BLCA dataset were downloaded to assess classifier robustness to different annotations and preprocessing pipelines. This included a version downloaded through the R package TCGABiolinks (27), a version from the Toil project (22), and one from the Recount3 project (28). Clinical data for the TCGA-BLCA cohort was downloaded from the Broad Institute (version 20160128, http://firebrowse.org/?cohort=BLCA). The IMVigor210 cohort was obtained through the IMvigor210CoreBiologies package (29) in TPM format. The classification was also evaluated on data preprocessed from the raw FASTQ files obtained from EGA (EGAD00001006960), but as the EGA identifiers were not linkable to the published metadata we only used the published RNA-sequencing data from IMvigor210CoreBiologies for this cohort. The UC-Genome (30) and UNC-108 (31) datasets were both obtained in processed TPM form from the authors. Raw CEL files were downloaded for microarray datasets Seiler2017 (GSE87304) (1) and Seiler2019 (GSE124305) (32). The Robertson T1 (3) was downloaded in FKPM and raw count format (GSE154261), and the Bowden T1 (33) was downloaded in FPKM format (GSE136401). The RotterdamBCG study cohorts A and B were obtained from the authors in FPKM format as used in the original study (34), as well as in Kallisto/Gencode39 preprocessed format. The UROMOL normalized counts were downloaded from the original publication (6). Collected datasets are described in Supplementary Table 2.

### Gene signatures used for validation

Predefined gene-signatures were applied to the data to validate the classification: early and late cell cycle genes, keratinization signature, *FGFR3* co-expressed genes, *TP63*, urothelial differentiation transcription factors (*PPARG, FOXA1, GATA3, ELF3*), urothelial differentiation genes (*UPK1A, UPK1B, UPK2, UPK3A, KRT20*), cell adhesion genes (*EPCAM, CDH1, CDH3*), MYC genes (*MYCL, MYCN, MYC*) and small-cell/neuroendocrine markers (*CHGA, SYP, ENO2*). Logtransformed and standardized values (Z-scores) were used to calculate several scores and ratios: the circuit score (35) to distinguish Uro from GU cases calculated as *RB1 + FGFR3 + CCND1 - E2F3 - CDKN2A* mRNA expression using log-transformed and standardized values (Z-scores), the Basal/Squamous ratio (36) calculated as *KRT5 + KRT14 - FOXA1 - GATA3*, the *ERRB* score as *EGFR - ERBB2 - ERBB3*, using standardized log2 mRNA expression values, and the late/early cell cycle ratio calculated as the median expression value of the late cell cycle signature genes minus the median expression value of the early cell cycle signature genes. Established cancer immune and stroma signatures used by the ESTIMATE tool (Immune141_ UP and Stromal141_ UP) were used (37). The ComplexHeatmap (v2.14.0) R package was used for heatmap visualization (38).

### Additional R packages

The R package ROCit (v2.1.1) was used for receiver operating characteristic (ROC) curves and area under the ROC curve (AUC) calculations. The R packages DistributionOptimization (v1.2.6) (39) and AdaptGauss (v1.5.6) (40) were used to fit Gaussian mixture models to the data and to determine cutoffs between distributions, respectively.

### Statistical tests

Fisher’s exact test was used to test the association of categorical variables (presence of mutations and response) with the molecular subtypes, and ANOVA was used for continuous variables. Bonferroni correction was used to correct for multiple testing. Kaplan-Meier analysis and log-rank test were used to visualize survival and compare survival outcomes.

### Addition of synthetic tumor microenvironment to the TCGA-BLCA cohort

We used 402 samples from the TCGA-BLCA cohort, quantified with Kallisto using the Toil Gencode23 index (22) and summarized into the filtered TPM format utilized by the Kassandra tumor microenvironment (TME) deconvolution study (19). We created 500 synthetic tumor microenvironment cell type compositions by generating random TME composition fractions between 0 and 1, multiplied with coefficients representing an average tumor microenvironment composition, comprising 18 cell types (19). The cell type fractions were rescaled to sum up to 1 for each of the 500 TME versions. For each synthetic TME, nine RNA-sequencing samples of each cell type were randomly selected from a cohort of 5677 RNA-sequenced samples in the same filtered Gencode23 TPM format. The average TPM expression profile of the 9 samples representing a cell type was rescaled to sum up to 106 and then multiplied by the precalculated cell type fraction and summed with the other cell types to form a pure synthetic TME TPM expression profile. Tumor and synthetic TME expression were summed together to generate 500 versions of the TCGA-BLCA dataset with sampled tumor fractions ranging from 0 to 1 (tumor TPM * tumor fraction + synthetic TME TPM * (1-tumor fraction)) (Supplementary Figure 1).

## Results

### Development of the LundTax2023 classifier

The Lund Taxonomy 2023 (LundTax2023) classifier was built utilizing a gene-pair rule-based RandomForest (RF) SSP using our R package multiclassPairs (23). Previous evaluations showed that this type of classifier enabled coherent bladder cancer subtype classification from array to RNA-sequencing data and vice versa, that high classification accuracy could be achieved in multiclass prediction of homogeneous data, and that training on data from multiple platforms was a feasible approach to reduce platform incompatibilities (10).

To create the new classifier, we first extended the classification of the Lund2017 study to our other datasets to form the training cohort. A five-class (Uro, GU, Ba/Sq, Mes-like, and Sc/NE) Lund Taxonomy RF-classifier was built using the well characterized Lund2017 microarray gene-expression dataset (n=301). We first identified subtype informative genes through RF within the Lund2017 dataset using the RMA preprocessed and Combat-batch corrected version of the dataset, without cohort scaling or gene centering, with data transformed into ranks (Figure 1B-1). To select sub-type informative features (genes), six models were trained, one predicting all classes, and one for each subtype in a 1-vs-rest configuration, using protein-coding genes with HGNC symbols present in both microarrays and RNA-sequencing datasets. From each model we selected the top 250 genes based on relative variable importance calculated by the RF model, resulting in a total of 1151 unique genes. A matrix of all possible binary gene-pair rule combinations (Gene X > Gene Y) was generated from raw RMA and SCAN data using the unique selected genes (Figure 1B-2). Six new RF-models were trained using this binary matrix as training data. The gene-pair rules were sorted by their variable importance according to each RF model, after which the top 500 rules were selected from each model, with a filter limiting the number of times a given gene was allowed to be used in a rule to 10. After removal of less contributing rules using the Boruta feature selection algorithm, the final 5-class model was trained using the 1127 retained rules (Figure 1B-3). We visualized the training cohort by a UMAP dimensionality reduction plot based on a Hamming distance matrix calculated from the binary gene-pair rules used by the classifier across all samples. The rules of the Lund2017 model separated subtypes into distinct clusters without notable platform separation of RMA and SCAN preprocessed data, but silhouette scores indicated some outlier samples where the UMAP cluster location was at odds with both the reference and out-of-bag (OOB) predicted subtype label (Supplementary Figure 2). A new model was produced excluding the 13 outlier samples, after which the outlier samples were reclassified (Figure 1B-4). The re-predictions were all in agreement with the outlier’s original cluster location. The outliers were reintroduced into the training set and used to build the starting model based on Lund2017. This model was applied to two additional uniformly preprocessed microarray datasets (Lund2020 and Lund2022) and 265 RNA-sequenced samples (s265) summarized to TPM using both Kallisto and Salmon (Figure 1C-1). All three datasets had in parallel also been classified by a set of 11 immunohistochemical markers on tissue microarrays (TMAs). Prediction results were in good agreement with both previous subtype classifications of these cohorts (83% concordance, average of 3 datasets), and with immunohistochemistry-based subtype assignments (72%). Classification concordance between RMA and SCAN versions of microarray data was 99% in Lund2020, 92% in Lund2022, and 99% between Kallisto and Salmon for the s265 RNA-sequencing dataset. The final predictor was built utilizing all four in-house datasets as training data, using the predicted labels on RMA and Kallisto data as final labels. Gene selection was performed for all four datasets using ranked Combat-adjusted RMA data for the microarrays and ranked Kallisto TPM data for the RNA-sequencing cohort (Figure 1C-2). Subtype-informative genes from all four datasets (n=717 genes) were combined into a gene-pair rule matrix using raw RMA, SCAN, Kallisto TPM, and Salmon TPM data for each sample (Figure 1C-3,4). A set of low-variance housekeeping genes (n=130), selected based on their stable expression across the four datasets, were also included to potentially act as stable pivot rule-partners to more high-variance subtype specific genes. Subsequently, a separate model was trained to classify Uro samples into UroA, UroB, and UroC. This was done by repeating the model training process on the subset of samples in the Lund2017 dataset that were classified as Uro both by the original labels and by the OOB predictions of the new classifier. This model was applied to the Uro classified samples of the other 3 datasets, after which a final model was trained on the UroA, UroB, and UroC samples of the entire cohort including all four datasets (Figure 1D).

**Fig. 2.**
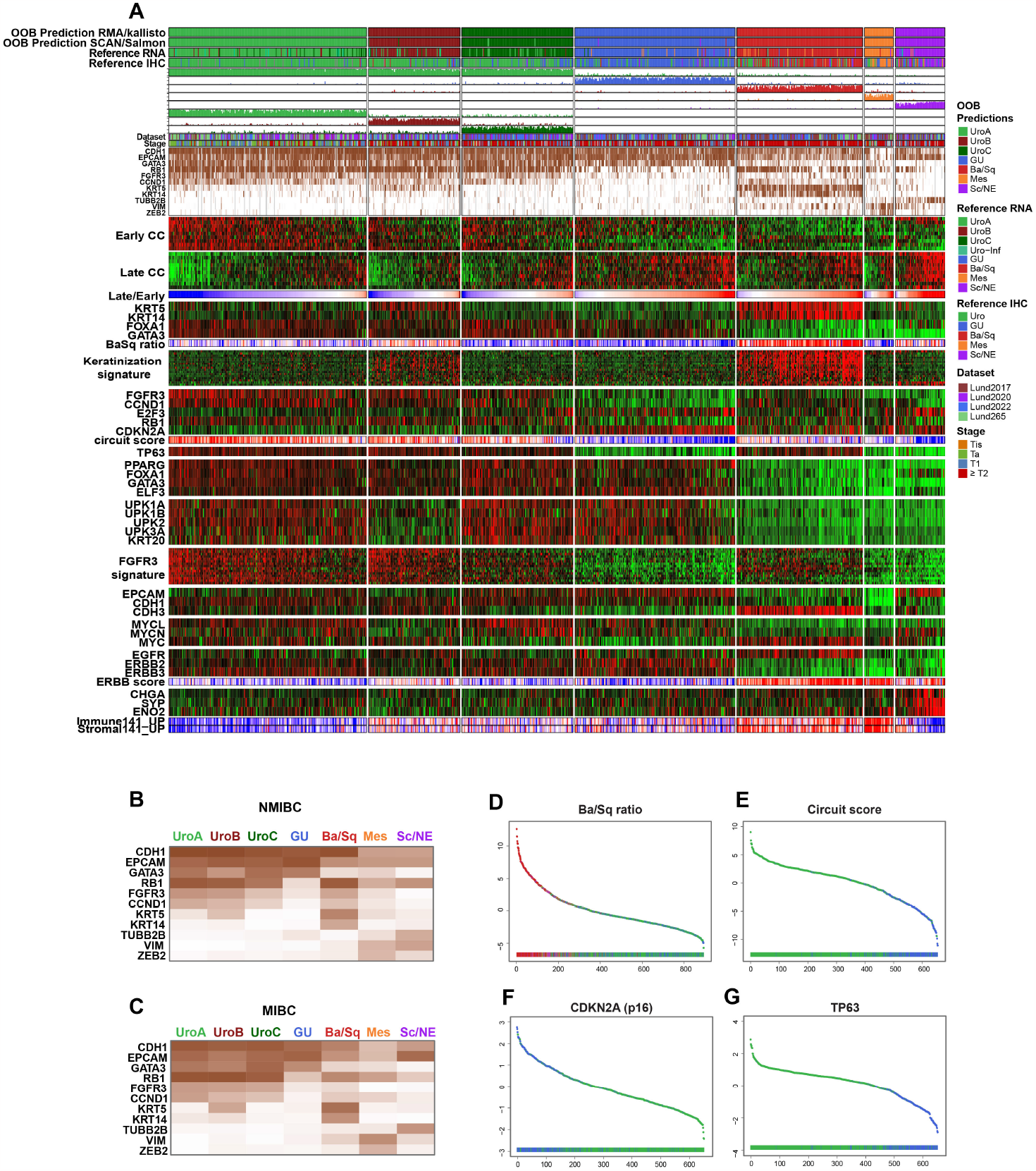
Final classification of the training data. **A:** The final classification and heatmap of the 894 samples included in the development of the LundTax2023 algorithm. From the top; out-of-bag (OOB) predictions for the RMA/Kallisto pre-processing version of the data; OOB predictions for the SCAN/Salmon pre-processing version; RNA reference; IHC reference; classification scores five-class system; classification scores for the Uro subclasses; dataset; pathological stage; intensity data for 11 IHC markers as indicated early cell cycle (Early CC) signature; late cell cycle (Late CC) signature; the ratio between late an early cell cycle ratio; gene expression levels for the four Ba/Sq defining markers; the calculated Ba/Sq ratio; keratinization gene expression signature; gene expression ratios for the genes distinguishing urothelial-like from genomically unstable tumors; the circuit score the expression levels of TP63; expression levels of urothelial differentiation transcription factors; expression levels of differentiation markers; expression of the FGFR3 gene expression profile; expression of cell-cell interaction genes; expression of the MYC family of transcription factors; expression of the EGF related receptor molecules EGFR, ERBB2 and ERBB3; the ERBB score; expression of neuronal marker gene specific for the Sc/NE subtype; immune infiltration score; stromal infiltration score. **B-C:** Average intensity levels of the specified IHC markers in the non-muscle invasive (**B**) and muscle invasive versions (**C**) of the molecular subtypes. **D-G:**Rank ordered plots showing samples ordered by Ba/Sq ratio (**D**), circuit scores (**E**), CDKN2A(p16) expression (**F**) or TP63 expression (**G**). Color code according to the five-class system. Color codes for gene scores and ratios: blue, low; red, high. IHC expression: white, low; brown, high.

**Fig. 3.**
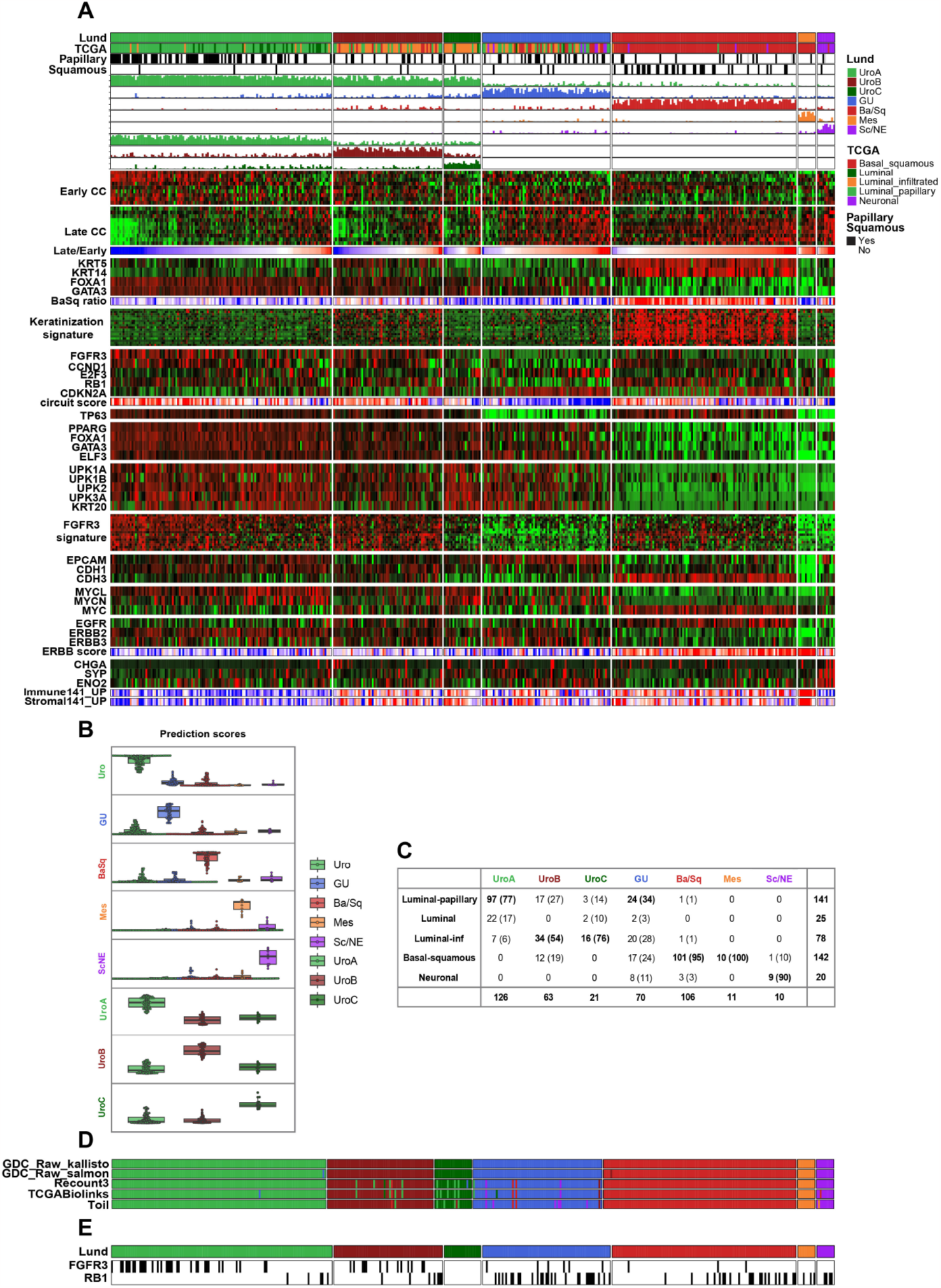
Classification results for the TCGA-BLCA cohort in the Kallisto TPM format. **A:** Gene expression heatmap of the TCGA-BLCA cohort. Classification scores and order of gene expression signatures as in Figure 2. **B:** Boxplots of classification scores for each molecular subtype. **C:** Subtype name translation table between the TCGA and LundTax2023 classes, n (%). **D:** Distribution of the Bonferroni corrected significant FGFR3 and RB1 mutations. Black, mutation; white, wild type. **E:** Classifications obtained using different preprocessing approaches.

### Biological evaluation of classification results by IHC and RNA signatures in the training cohorts

Expression signatures for the four in-house datasets were visualized together using Z-scores calculated for each dataset separately (Figure 2A) using RMA and Kallisto preprocessed data. OOB predictions on the four training sets were highly concordant between the RMA and Kallisto data and the alternative SCAN and Salmon preprocessing (Figure 2A). The OOB prediction scores were also similar between preprocessing versions (Supplementary Figure 3).

IHC markers CDH1, EPCAM, GATA3, RB1, FGFR3, CCND1, KRT5, KRT14, TUBB2B, VIM, and ZEB2, scored on tumor cells, were well aligned with the RNA-based classification. CDH1 and EPCAM were detected across most samples but lower in the Mes-like and Sc/NE subtypes that deviated most from an epithelial cell state. RB1 and CCND1 were predominantly lower in GU and Sc/NE tumors. FGFR3 was predominantly seen in Uro-classified tumors, while GATA3 expression encompassed the “luminal” tumors, also including GU. KRT5 expression was strongly positive across most Ba/Sq classified tumors and to a lower extent in UroB tumors, while KRT14 expression was confined largely to Ba/Sq tumors. TUBB2B expression was seen mainly in Sc/NE classified samples, while tumor cell expression of VIM was mainly seen in Mes-like samples and some Ba/Sq and Sc/NE tumors. Similarly, ZEB2 expression was predominantly confined to Mes-like classified samples. In Figures 2B and 2C we have separated non-muscle invasive samples of UroA, UroB, UroC, and GU from their respective muscle invasive versions and show that there are no essential differences of cancer cell expression levels among the reported markers.

Cross-cohort expression signature patterns reflected previous observations (Figure 2A). Briefly, Uro tumors showed expression patterns indicating lower proliferation, with high expression of early cell cycle genes and low expression of late cell cycle genes compared to other subtypes. The per sample ratio of these cell cycle signatures was used to order tumors within each subtype and was overall lower in the Uro subtype. The consensus definition of Ba/Sq tumors, with high expression of basal markers *KRT5* and *KRT14* and low expression of luminal marker *FOXA1* and *GATA3* clearly overlapped with Ba/Sq classification results. UroB tumors showed KRT5 expression, but not uniform *KRT14* expression, and both Mes-like and Sc/NE tumors displayed low expression of all four genes. A rank-order version of the Ba/Sq ratio data is presented in Figure 2D). Broad expression of keratinization related genes was confined to Ba/Sq tumors and in a less consistent manner in UroB tumors. The genomic circuit score (see) used to delineate GU tumors from Uro samples (35) was low in GU tumors, indicating that features associated with this group were recapitulated, including reduced *FGFR3* and *CCND1* expression, elevated *E2F3* expression, loss of *RB1* expression, and retained *CDKN2A* expression. A rank-order version of the circuit score ratio data is presented in Figure 2E and a separate rank ordered expression values for *CDKN2A* in Figure 2F. Reduced expression of *TP63* further demarcated GU from Uro tumors (Figure 2G). Despite using combined Z-scores from each separate dataset without further batch adjustments, expression rank plots of the Ba/Sq ratio, circuit score, *CDKN2A*, and *TP63* separated the subtypes well. Both transcription factors associated with differentiation and markers of terminal differentiation were more highly expressed across Uro and GU tumors compared to the non-luminal subtypes, however within the luminal category the UroB samples showed the lowest expression of terminal differentiation markers such as UPKs and *KRT20*. Expression of a *FGFR3* signature was strongest in UroA and UroB, while UroC and Ba/Sq showed a lower and more sporadic expression pattern of the genes of this signature. Uro cases also show higher expression of the adhesion genes *EPCAM* and *CDH1*, and lower expression of *CDH3*. This pattern shifts in the Ba/Sq samples, instead showing high expression of *CDH3*. The *MYC* family of transcription factors shows previously identified expression patterns, with high expression of *MYCL* and *MYCN* but low expression of *MYC* in Uro, especially in UroC, and in GU samples, as opposed to Ba/Sq cases, whereas UroB samples display a mixed expression of these factors. Expression levels of epidermal growth factors *ERBB2* and *EGFR* also conformed with earlier Lund Taxonomy results, with elevated *ERBB2* and *ERBB3* expression and lower expression of *EGFR* in GU samples with the reverse expression pattern in Ba/Sq tumors. Expression of the neuroendocrine lineage markers *CHGA, SYP*, and *ENO2* was seen primarily in Sc/NE classified tumors. Taken together, the expanded cohort obtained by transferring the Lund2017 classification to additional array and RNA-sequencing datasets and subsequently utilized to train the new optimized and more comprehensive LundTax2023 model, displays highly coherent expression signatures and features of the different molecular subtypes at both the RNA and IHC levels.

### Applications to independent MIBC datasets

The Lund-Tax2023 classifier was applied to several datasets to determine whether characteristic subtype-associated gene signatures and other properties were coherently recapitulated in independent cohorts and datasets.

#### TCGA-BLCA

We applied the LundTax2023 to the TCGA-BLCA cohort (2), using 407 MIBC tumors in Kallisto TPM data format. The prediction resulted in 209 Uro cases (51%), 73 GU (18%), 105 Ba/Sq (26%), 10 Mes-like (2.5%), and 10 Sc/NE (2.5%) cases (Figure 3A). The Urothelial-like subclassification produced 126 UroA (60% of all 209 Uro-classified samples), 62 UroB (30%) and 21 UroC (10%) cases, respectively. The prediction scores were distinct (ANOVA, p *<* 10^−16^), with few ambiguities. (Figure 3B). The samples predicted as Ba/Sq showed high Ba/Sq scores (*KRT5, KRT14*/*FOXA1, GATA3*) and high expression of keratinization gene signature. The Ba/Sq cases also showed a coherent pattern of elevated *MYC* expression and low *MYCL* and *MYCN*. The circuit score distinguished the GU cases tumors from the Uro, indicating that GU was correctly classified, strengthened by the characteristic loss of *TP63* expression in the GU subtype. The expression of luminal markers mirrored the training set, with transcription factors and UPKs and *KRT20* expression elevated across Uro and GU. Within the luminal subtypes the *FGFR3* gene signature was more pronounced in UroA and UroB compared to UroC and GU. The Sc/NE was negative for *KRT5, KRT14, FOXA1*, and *GATA3* expression, negative for the urothelial differentiation markers (*PPARG, FOXA1, GATA3*, and *ELF3*), as well as for the *FGFR3* gene expression signature, but expressed neuroendocrine marker genes *CHGA, SYP*, and *ENO2*, and had high early/late cell cycle ratios. The Mes-like subtype was more or less negative for most of the applied gene signatures but showed the highest scores among all subtypes for Immune141_UP and Stromal141_UP, indicating a high level of infiltration of non-tumor cells. From this, we conclude that the LundTax2023 algorithm classifies the TCGA dataset according to the LundTax system into distinct and biologically coherent groups. We compared the LundTax 7-group classification with the TCGA classification (Figure 3C). UroA corresponded to TCGA Luminal subtypes, and UroB both to Luminal subtypes and to TCGA basal-squamous. UroC mainly corresponded to Luminal infiltrated, and GU to Luminal. However, a fairly large proportion of GU was also classified as TCGA Basal-squamous and Neuronal. Almost all LundTax Ba/Sq corresponded to TCGA Basal-squamous, as well as almost all Sc/NE to TCGA Neuronal. All Mes-like were classified as TCGA Basal-squamous. Hence, even though the LundTax Urothelial-like nearly exclusively overlapped with TCGA Luminal subtypes they differ radically with respect to the internal divisions. No equivalent to GU is present in the TCGA system. The TCGA Basal-squamous is more inclusive than the LundTax Ba/Sq, including UroB and Ba/Sq, as well as Mes-like cases. The TCGA Neuronal is also more inclusive compared to LundTax Sc/NE subtype. We then applied the LundTax2023 algorithm to the Salmon TPM preprocessed and three alternative external preprocessing versions of the TCGA-BLCA dataset, (Recount3, TCGABiolinks and Toil). In the 7-class solution, 2 cases from the Salmon version, 17 cases from Recount3, 20 cases from TCGABiolinks, and 17 cases from Toil received discordant predictions compared to the raw data, equivalent to an overall 1-5% discordance (Figure 3D). For the three independently preprocessed versions, almost half of the discrepancies occurred between the Uro subclasses (9/17 for Recount3, 10/20 for Biolinks, and 6/17 for Toil), resulting in an overall 2-3% discordance when comparing the 5-class solutions. We used the reported gene mutation data and used Fisher’s exact tests with Bonferroni correction to identify mutations associated with the LundTax subtypes. FGFR3 mutations were enriched in the UroA and UroB groups, but absent in UroC, whereas RB1 mutations were significantly enriched in GU (p *<* 10^−5^). The LundTax UroC showed absence of both FGFR3 and RB1 mutations (3E). From this we conclude that the algorithm is quite insensitive to different preprocessing methodologies and separates subtype-associated genomic alterations.

#### IMVigor210

We applied the LundTax2023 to the IMVigor210 cohort with 347 samples from patients with locally advanced or metastatic tumors treated with checkpoint inhibition (CPI) using TPM values obtained from the IMvigor210CoreBiologies R package (29). A total of 176 samples were classified as Uro (50%), 64 as GU (18%), 90 as Ba/Sq (26%), 9 as Mes-like (3%) and 9 as Sc/NE (3%), with the Uro group subdivided into 71 UroA (40% of Uro), 70 UroB (40%) and 35 UroC (20%) cases (Figure 4A). Prediction scores were again very distinct (Supplementary Figure 4). The Ba/Sq ratio, keratinization signature, EGFR relative ERBB2 expression, and *MYC*-gene expression clearly delineated the Ba/Sq group. The luminal tumors Uro and GU both showed expression of the differentiation-related transcription factors, as well as several of the differentiation markers, with the circuit score and *TP63* expression distinguishing Uro from GU. Again, the Mes-like tumors showed low expression of almost all the applied gene signatures, whereas Sc/NE samples showed strong expression of *EPCAM* and the neuroendocrine specific markers. From this we conclude that the algorithm classifies the IMVigor210 data into distinct and biologically coherent LundTax subtypes. Mutation data for 392 genes was available in 274 samples, with *FGFR3, RB1*, and *TP53* gene mutations significantly associated with the subtypes (Figure 4B). Thus, *FGFR3* mutations were frequent in UroA (52%) and UroB (27%), while almost absent from UroC (7%) and GU (4%). *RB1* mutations were significantly enriched in GU (p *<* 10^−5^), but also frequent in Sc/NE. *TP53* mutations were more common in GU, UroC, Sc/NE, and Ba/Sq whereas UroA and UroB showed lower frequencies. We then made use of the immune phenotype mapping provided in the IMVigor210 dataset, in which the immune infiltration pattern was characterized as either desert, excluded, or inflamed (41) (Figure 4C). The least infiltrated group was UroA, with a decreasing fraction of desert-cases in UroB, UroC, GU and Ba/Sq tumors, respectively. The fraction of infiltrated tumors remained the same going from UroA to Ba/Sq whereas the fraction of inflamed increased in this direction. The Mes-like subtype was the most inflamed, whereas Sc/NE tumors displayed a predominance of the excluded infiltration pattern. This shows that in addition to being well defined at the cancer cell phenotype level, the LundTax subtypes also show distinct infiltration patterns. We used the reported atezolizumab response data and noted that the subtype with the best response, defined as partial or complete response, was Sc/NE, followed by GU, whereas the response rate was very low in Ba/Sq, and absent in nine patients with Mes-like tumors (Figure 4D). No complete responder was observed among 71 patients with UroA-tumors.

**Fig. 4.**
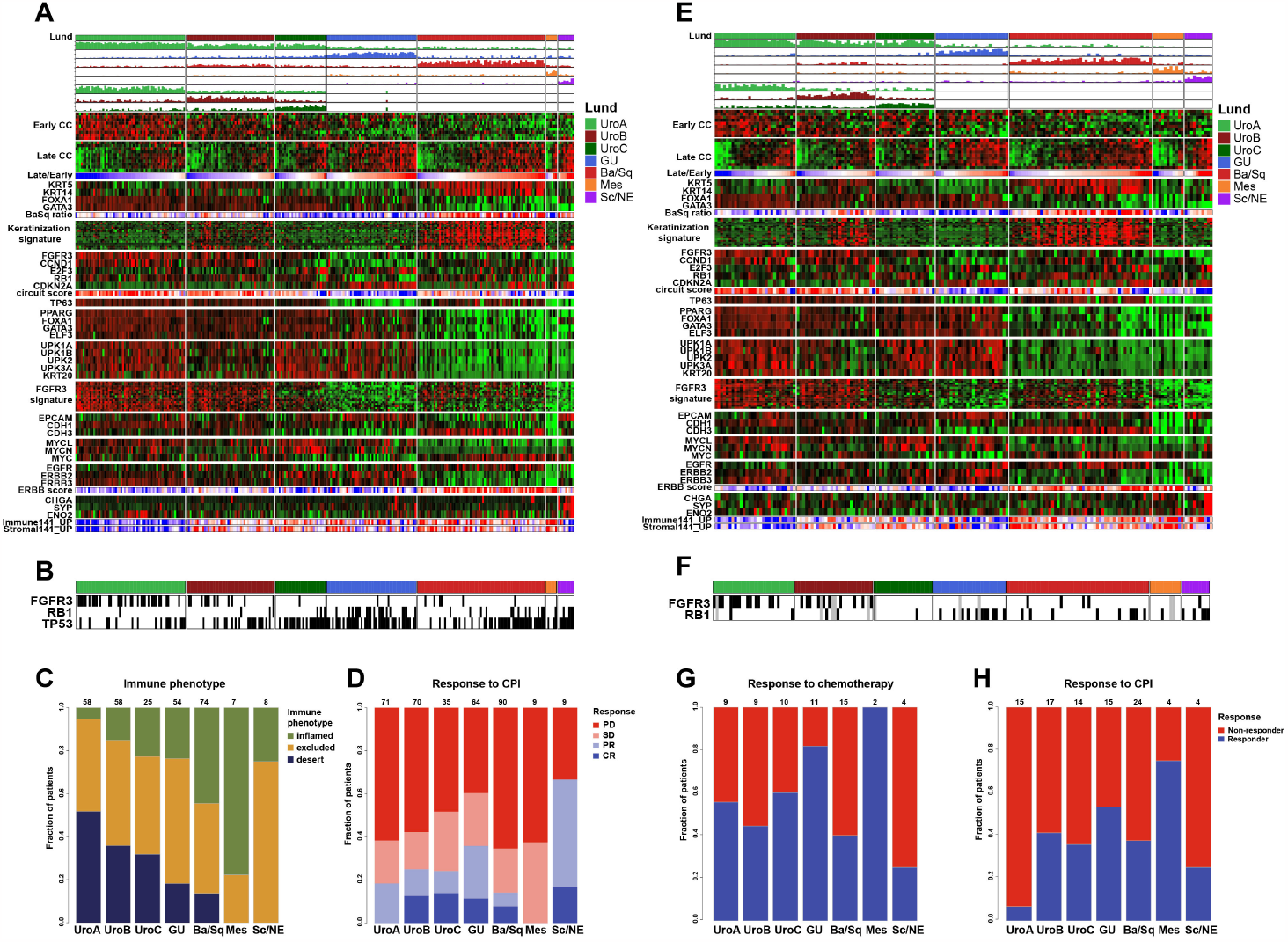
Classification results for the IMVigor210 and UC-genome cohorts. **A:** Gene expression heatmap of the IMVigor210 cohort. Classification scores and order of gene expression signatures as in Figure 2. **B:** Distribution of the significant gene mutations after Bonferroni correction: FGFR3, RB1 and TP53. Black, mutation; white, wild type; gray, no data. **C:** Distribution of the immune infiltration patterns “desert” “excluded”, and “inflamed” across the molecular subtypes. **D:** Distribution of the CPI response pattern across the molecular subtypes. **E:** Gene expression heatmap of the UC-genome cohort. Classification scores and order of gene expression signatures as in Figure 2. **F:** Distribution of the Bonferroni corrected significant FGFR3 and RB1 mutations. Black, mutation; white, wild type; gray, no data. **G-H:** Distribution of the chemotherapy (**G**) and CPI (**H**) response across the molecular subtypes.

#### UC-Genome

The UC-Genome cohort consists of 218 samples from patients with metastatic urothelial carcinoma (30) RNA-sequencing data for 176 samples was available. LundTax classification resulted in 78 Uro, (44%), 26 GU, (15%), 51 Ba/Sq, (29%), 11 Mes-like (6%), and 10 Sc/NE cases (6%). Subclassification of Uro produced 29 UroA (37%), 28 UroB (36%) and 21 UroC (27%) cases (Figure 4E), again with distinct prediction scores (Supplementary Figure 4). Classifications were validated by the applied gene expression scores and signatures (Figure 4E). *FGFR3* mutations were almost exclusively detected in UroA and UroB, and again absent in UroC and GU cases (Figure 4F). Using the provided response data to systemic chemotherapy and CPI, we observed that GU responded the best and Ba/Sq the worst to chemotherapy, in line with recent results (42), whereas the Urothelial-like subtypes (UroA, B and C) were intermediate (Figure 4G). We again noticed that UroA was largely refractory to CPI treatment (Figure 4H).

#### UNC-108

UNC-108. The UNC-108 cohort (31) consisting of 89 samples from patients with metastatic UC treated with immune checkpoint blockade was classified into 43 Uro (48%), 16 GU (18%), 23 Ba/Sq (26%), 2 Mes (2%) and 5 Sc/NE (6%) tumors. Within the Urothelial-like subtype, 23 samples were classified as UroA (53%), 14 as UroB (33%) and 6 as UroC (14%). Prediction scores were again distinct. The applied gene expression scores and signatures validated the classification. FGFR3 gene mutations were significantly associated with UroA and UroB (p *<* 10^−3^) and absent in UroC and GU. This study also provided response data to CPI. Again, UroA tumors showed almost no response to CPI treatment (Supplementary Figure 5).

#### Seiler2017

The Seiler2017 neoadjuvant chemotherapy (NAC) cohort consists of 305 MIBC samples analyzed on the Affymetrix Human Exon 1.0 ST Array and subtyped according to the Genomic Subtyping Classifier (GSC) (1). We generated two preprocessing versions of this dataset, RMA and SCAN. For the first version (RMA normalization), the LundTax2023 algorithm predicted 142 Uro (47%), 62 GU (20%), 88 Ba/Sq (29%), 5 Mes-like (2%), and 8 Sc/NE (3%) samples (Figure 5A). Within the Uro group, we obtained 58 UroA (19%), 27 UroB (9%) and 54 UroC (19%) cases. When comparing prediction results between the RMA and SCAN preprocessing pipelines we obtain a concordance of 93.4% (285/305) and 90.8% (277/305) for the 5-class and 7-class solution, respectively. The most frequent discrepancies were in samples classified as GU in the SCAN version, 71 cases, of which 7 were classified as Uro in the RMA version, both subtypes belonging to the luminal class of tumors. Prediction scores were distinct (Figure 5B), and classifications were validated by the applied gene expression scores and signatures (Figure 5A).

**Fig. 5.**
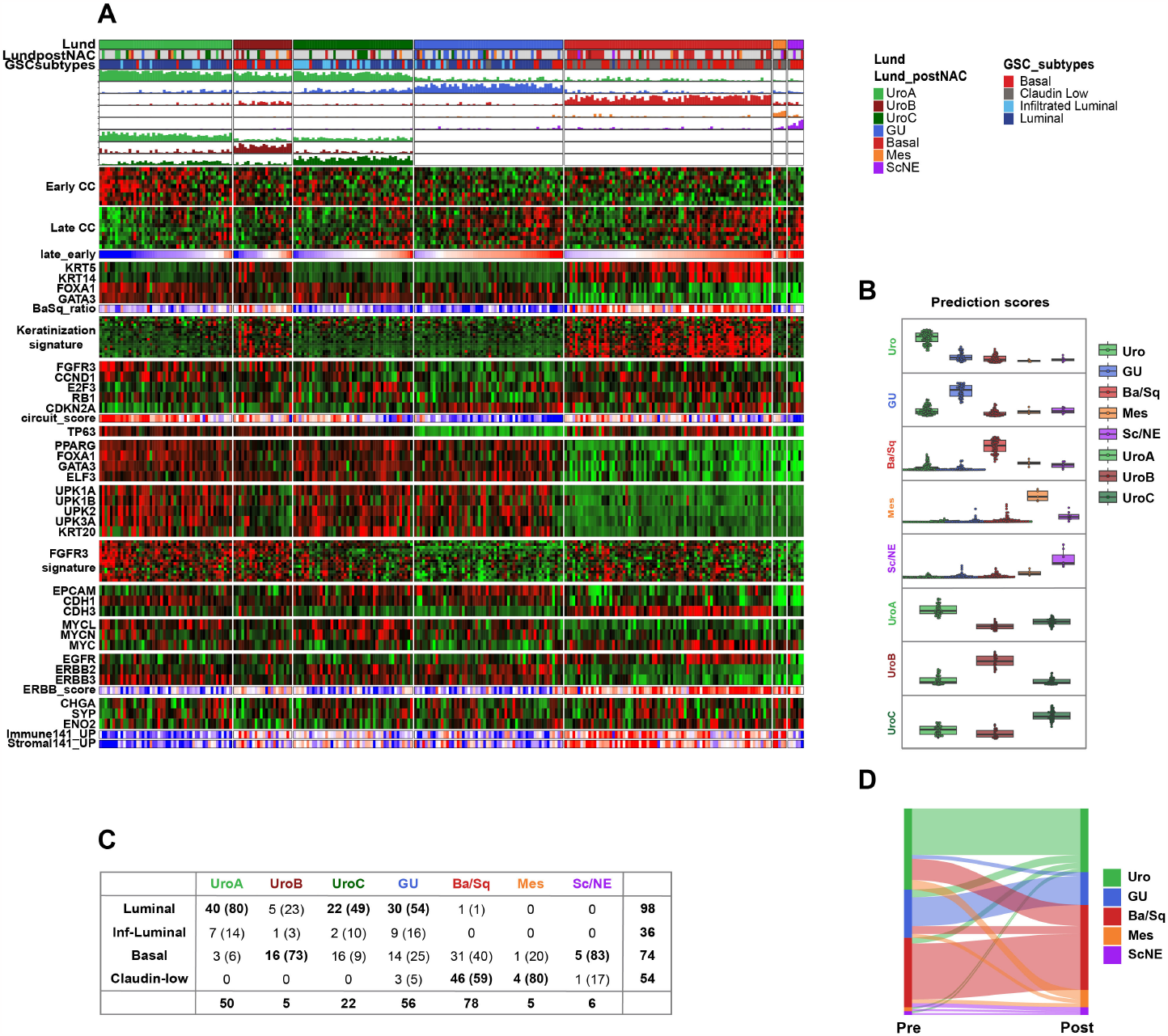
Classification of the Seiler2017 microarray-based cohort. **A:** Gene expression heatmap of the Seiler2017 cohort. Classification scores and order of gene expression signatures as in Figure 2. **B:** Boxplots of classification scores for each molecular subtype. **C:** Subtype name translation table between the GSC and LundTax2023 classes, n (%). **D:** Sankey diagram showing subtype shifts between pre- and post-chemotherapy samples according to LundTax2023 classification.

A comparison between the LundTax and GSC classification schemes (Figure 5C) showed that Uro mainly corresponded to the GSC Luminal class, but also included GSC Infiltrated Luminal and Basal classes. A similar pattern was seen for the LundTax GU. The LundTax Mes-like group was found mainly in the Claudin Low group, and Sc/NE in the Basal group. Of the 23 Urothelial-like classified as Basal according to GSC, 16 were of the UroB subtype, known to show features resembling Ba/Sq (11). Of the GSC Infiltrated luminal group, 27 samples were classified as LundTax Uro, but dominated by UroC (19/27), and 9 as GU. We then compared the subtype prediction of 107 samples with matched post-NAC samples provided in (32). The authors reported that half of the samples (52.6%) change subtype, with 40% of the luminal samples acquiring a Basal subtype after treatment. When we apply the 5-class solution to matched samples we observe that 34.5% of the samples changed subtype, decreasing to 27.1% if changes involving the Mes-like subtype were disregarded, as they may represent biopsies with a very low tumor cell content.

### Classification and low purity

To evaluate how tumor purity affects classification we made use of the Kassandra tumor microenvironment (TME) composition prediction work (19). We preprocessed 402 samples from the TCGA-BLCA cohort following the same pipeline used in the Kassandra study and we created 500 synthetic TME compositions that were added to the original TCGA-BLCA with TME fractions ranging between 0 and 1, creating 500 versions of the TCGA-BLCA dataset of varying purity. The LundTax2023 classifier and the reference consensus MIBC classifier (5) were then applied to each of the 500 composite datasets to examine the effect on classification results. As tumor biopsies may vary in purity, we limited the analyses to cases that originally showed a purity exceeding 70% tumor cell fraction as assessed by the Kassandra TME tool, resulting in 252 cases (Figure 6A). With increasing infiltration, Uro samples gradually altered classification towards GU and Ba/Sq. Past approximately 60% added synthetic TME content, Uro, GU, and Ba/Sq predictions shifted towards Mes-like. Mes-like samples all retained their Mes-like classification, while Sc/NE samples underwent a more rapid shift towards Mes-like classification. The consensus MIBC predictor deviated more from the original predictions and did so at a lower fraction of added TME (Figure 6B). While the LundTax2023 predictions drifted towards Mes-like, the overall pattern for the consensus MIBC predictions was more erratic, with an initial shift towards Stroma-rich, and then a consistent shift towards Ba/Sq. The behavior of both classifiers was consistent also when looking at the full cohort of 402 samples (Supplementary Figure 1).

**Fig. 6.**
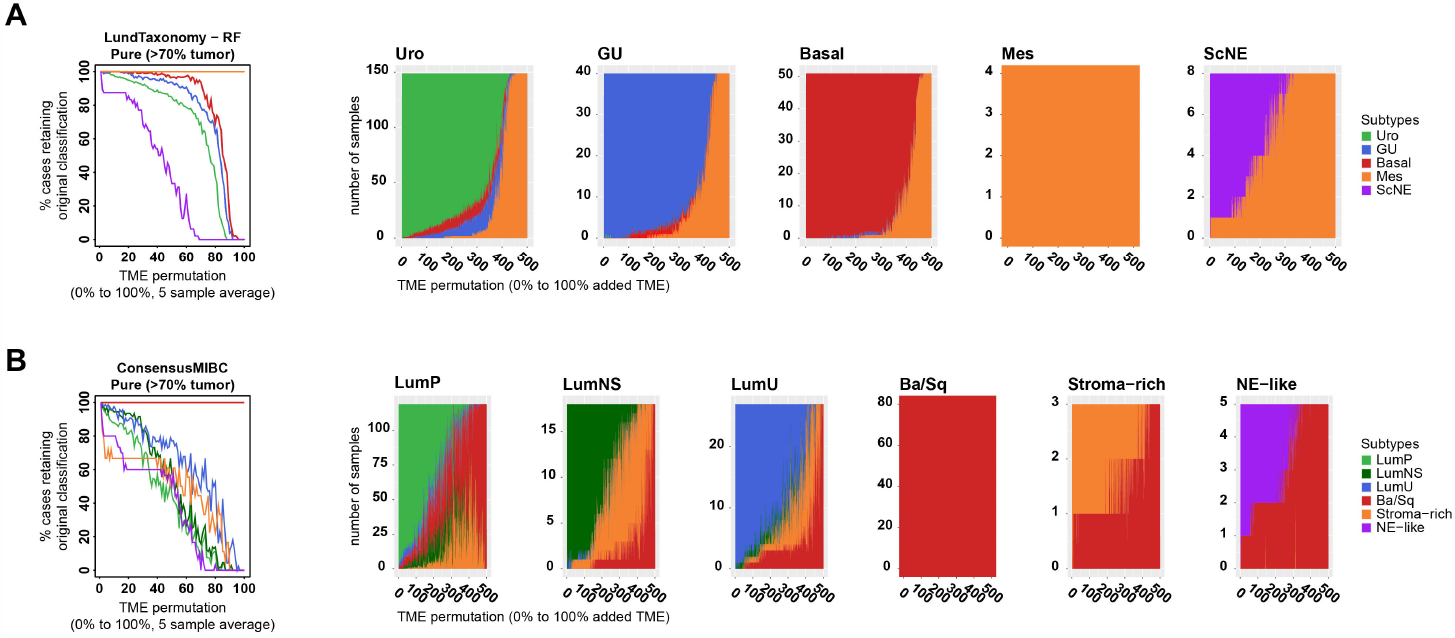
Effect of tumor purity on classification stability. Cohort of 500 synthetic TME compositions created in silico increasing the level of infiltration in a stepwise manner. The data was classified using the LundTax2023 (**A**) or the consensus classification (**B**).

When Uro predicted cases were divided into UroA, UroB, and UroC classification shifts towards Ba/Sq were almost entirely confined to UroB cases, while UroC saw shifts more often towards GU compared to tumors subclassified as UroA.

### Summary from MIBC cohorts

From the presented results we conclude that the updated and optimized version of the LundTax classifier accurately identifies established molecular subtypes regardless of cohorts, preprocessing algorithms, and platforms. When the proportions of the predicted subtypes across the different datasets are compared (Figure 7A), the subtypes maintain similar proportions across cohorts. The proportions among the Uro subdivisions varied slightly more, with UroA comprising 34-60% of the total Uro cases, UroB 29-40%, and UroC 10-40% among all samples in all datasets. The distribution of FGFR3 and RB1 mutations is also consistent and significantly correlated with subtypes across cohorts (Figure 7B). FGFR3 mutations are frequent in UroA and UroB, present in 33 and 24% of the cases, respectively, but almost absent in UroC (2.7%) and GU (5%), while the GU subtype is enriched for RB1 mutations, present in 39% of the cases. We also conclude that the LundTax2023 version is less influenced by low sample purity compared to the reference consensus MIBC classifier.

**Fig. 7.**
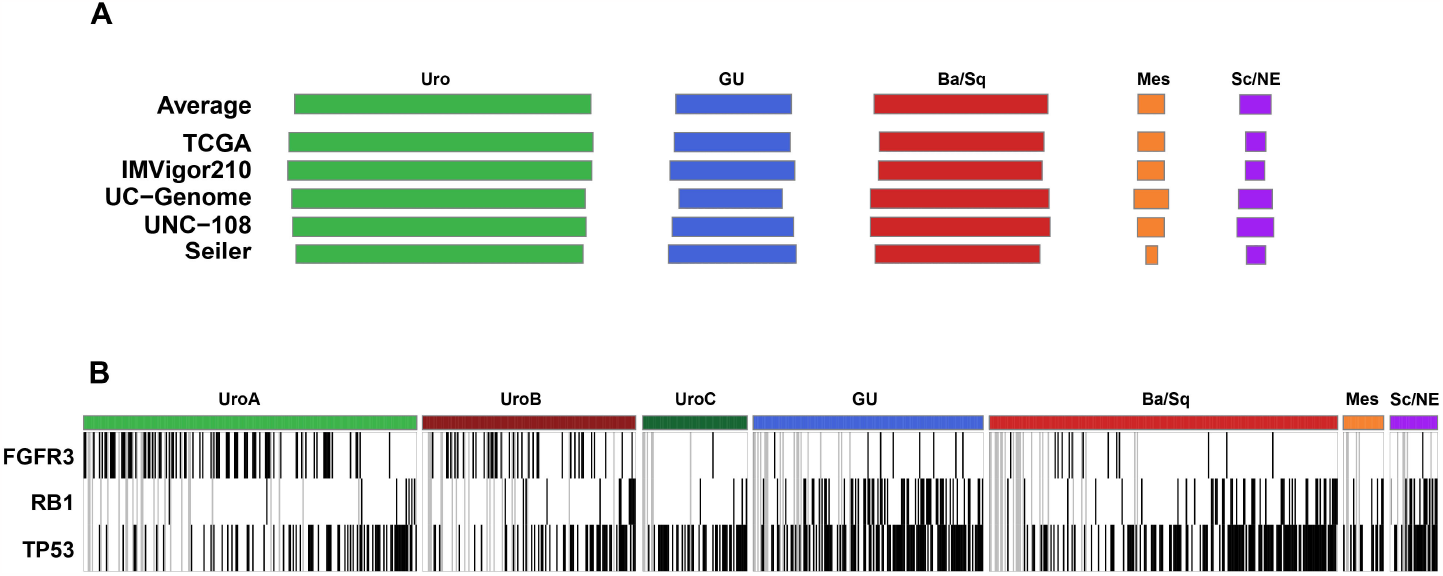
A summary of classification and mutation data from the muscle invasive cases. **A:** The variation of the five-class proportions in the individual five muscle invasive cohorts. **B:** The pooled mutation data for FGFR3, RB1, and TP53 mutations. Black, mutation; white, wild type; gray, no data.

### Application to external and independent NMIBC datasets

#### UROMOL and Leeds

We have previously analyzed the UROMOL and Leeds cohorts using an earlier version of the classification algorithm (43). The UROMOL cohort includes 535 NMIBC samples, with a majority of Ta cases (397/535). We applied the LundTax2023 classifier and obtained 510 Uro cases (95%), 22 GU (4%), 1 Ba/Sq (0.2%) and 2 Mes-like (0.4%). Subclassification of Uro samples resulted in 456 UroA (89%), 43 UroB (8%) and 11 UroC cases (2%). The Leeds dataset is composed of 217 samples, including Ta (113) and T1 (104) cases. The classification resulted in 203 Uro cases (93%), 12 GU (6%) and 2 Ba/Sq (1%), with Uro samples subclassified into 191 UroA (94%), 11 UroB (5%) and 1 UroC (0.5%). The concordance between the previous and the present LundTax2023 classifications were 99% for both cohorts using the 5-class solution, and 99% and 90% for UROMOL and the Leeds cohorts using the 7-class solution, respectively (Supplementary Figure 6).

#### RotterdamBCG

The cohort consists of two datasets of 132 (cohort A) and 150 (cohort B) cases, respectively, with high grade (WHO 2022) non-muscle-invasive tumors treated with adjuvant BCG-instillations (34), stratified into one out of three defined BCG response subtypes (BRS 1-3). The cohort shows an extremely high progression rate of 34% (95/282) compared to the current 6-8% progression rate at five years in a Swedish population-based registry (44). We applied the LundTax2023 classifier and obtained 208 Uro (74%), 55 GU (20%), 10 Ba/Sq (4%), 2 Mes-like (0.7%), and 4 Sc/NE (1%) predicted cases, and within the Uro group, 166 UroA (79%), 20 UroB (10%) and 22 UroC (11%) (Figure 8A). The prediction scores were distinct (Figure 8B), and classifications were validated by the gene expression ratios and signatures (Figure 8A). We noticed that BRS3 cases showed high levels of infiltration assessed by immune and stromal infiltration scores (Figure 8A). We therefore produced a generalized infiltration score for each case using the Immune141_UP and Stroma141_UP average Z-values. This score predicted the BRS3 subtype with an AUC of 0.86 (Figure 8C), reiterating that lower cancer cell purity is a characteristic feature of the BRS3 tumors (34). According to the authors, 59% (20/34) of the BCG treated tumors change BRS subtype after treatment, with the majority (14/20) converting to the BRS3 subtype. However, using the LundTax2023 classification only 12% (4/34) changed subtype after treatment (Figure 8D). Hence, our interpretation is that tumors that recur/progress after BCG treatment do not change cancer cell phenotypes but rather show a lower level of cancer cell purity, i.e., a higher level of infiltration.

**Fig. 8.**
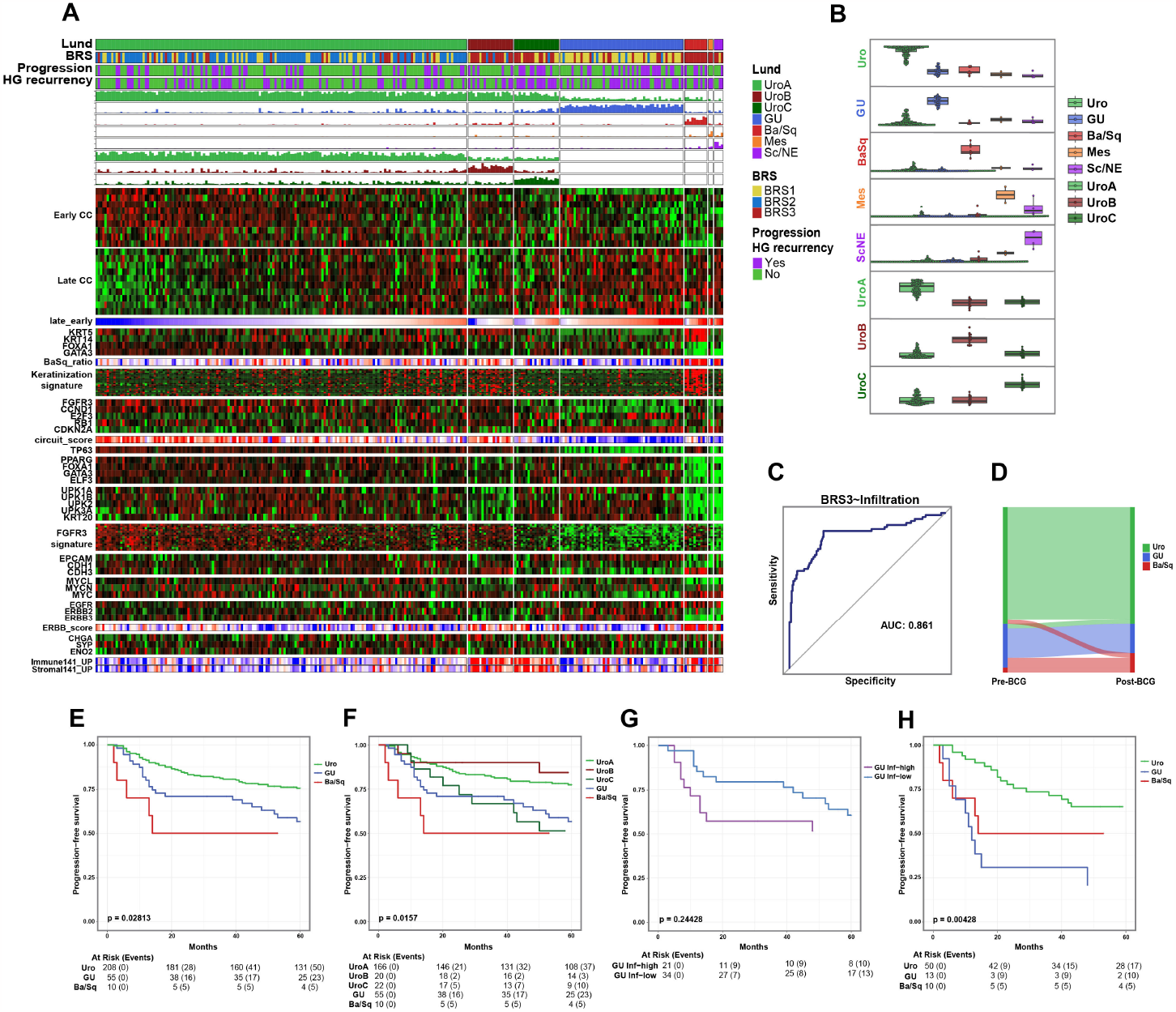
Classification of the Rotterdam cohort of high grade T1 tumors (cohort A+B, n = 279). **A:** Gene expression heatmap of the Rotterdam cohort. Classification scores and order of gene expression signatures as in Figure 2.**B:** Boxplots of classification scores for each molecular subtype. **C:** Prediction of the BRS3 subtype versus BRS1/2 using a merged immune and stromal score. **D:** Sankey diagram showing subtype shifts in 34 patients with matched pre- and post-BCG tumor samples, according to LundTax2023 classification. **E-H:** Kaplan-Meier plots using progression as endpoint. Samples stratified according to the five-class (**E**) or seven-class (**F**) LundTax system. GU samples were separated into high and low infiltrated cases (**G**) and the pooled BRS3 cases were analyzed using the five-class LundTax2023 (**H**). The Mes-like (n = 2) and Sc/NE (n = 4) groups were excluded from the analysis.

We then used the 5- and 7-class LundTax2023 classifications to test the association between subtype and progression after BCG treatment. Only Uro, GU and Ba/Sq cases were included in the 5-class Kaplan-Meier analysis as there were too few Mes-like and Sc/NE cases (Figure 8E). The Uro tumors showed the best PFS and Ba/Sq the worst, as previously reported in population-based data with primary T1 tumors (45). In the 7-class solution, the Ba/Sq group still showed the worst PFS, and UroA and UroB fared the best (Figure 8F). We then used the previously calculated infiltration score to divide the GU samples into high and low infiltration groups using a ROC analysis optimized threshold. This analysis revealed that highly infiltrated GUs had as bad prognosis as the Ba/Sq (Figure 8E and G). Hence, by combining LundTax cancer cell phenotype with an independent index for infiltration, an adjusted estimate of PFS can be obtained. We then classified all the BRS3 cases (n=76) from the combined datasets according to the LundTax system. The Kaplan-Meier curves showing PFS suggested that the BRS3 group is heterogeneous both with respect to LundTax molecular subtypes as well as for progression risk (Figure 8H).

#### RobertsonT1

The Robertson T1 cohort is comprised of 73 primary T1 tumors with RNA extracted from FFPE material and sequenced using 50bp single-end reads (3). The LundTax2023 classification of this cohort yielded 63 Uro (86%) and 10 GU (14%) predictions (Figure 9A). The further sub-classification of the Uro group produced 55 UroA (87%), 7 UroB (11%), and 1 UroC (2%). The prediction scores were unambiguous. To clarify the relationship between the Robertson classification and the LundTax classification, we first plotted the expression level of the *TP63* gene in the respective Robertson classes (Figure 9B). This clearly identified the Robertson T1-LumGU as GU. We then applied the keratinization signature, expected to be high in UroB, which identified the Robertson T1-Inflam group as the LundTax UroB equivalent (Figure 9C). UroB showed a higher level of infiltration compared to the rest of the luminal subtypes, as determined by the Immune141_UP and Stromal141_UP scores (Figure 9A). The remaining three Robertson classes showed higher expression of the *FGFR3* signature than T1-LumGU (GU), and at similar levels as in the T1-Inflam/UroB, indicative of a UroA profile (Figure 9D). Robertson states that the T1-TLum subtype shows repressed proliferation hallmarks. To investigate the existence of a natural threshold between low and high proliferative UroA tumors, we analyzed the distribution of the proliferation index using a Gaussian Mixture Model (GMM) that indeed indicated a low and high proliferation group of UroA tumors (Figure 9E). We then investigated the expression of the luminal differentiation gene *RXRA*, associated with the T1-TLum group, and observed that both the T1-TLum and the low-proliferation UroA tumors showed a higher *RXRA* expression. According to Robertson et al., the T1-Early and T1-Myc subtypes were associated with elevated Myc activity. *MYC* expression across subtypes of both classification systems (Figure 9F) showed that although the T1-Early and T1-Myc subtypes had elevated *MYC* expression, T1-TLum and T1-TInflam also had relatively high expression with T1-LumGU/GU tumors being the only group with low *MYC*. Taken together, we show that the Robertson’s T1 molecular subtypes demonstrate a good concordance with the LundTax subtypes when adding the proliferation variable, not considered a classification feature by the LundTax system.

**Fig. 9.**
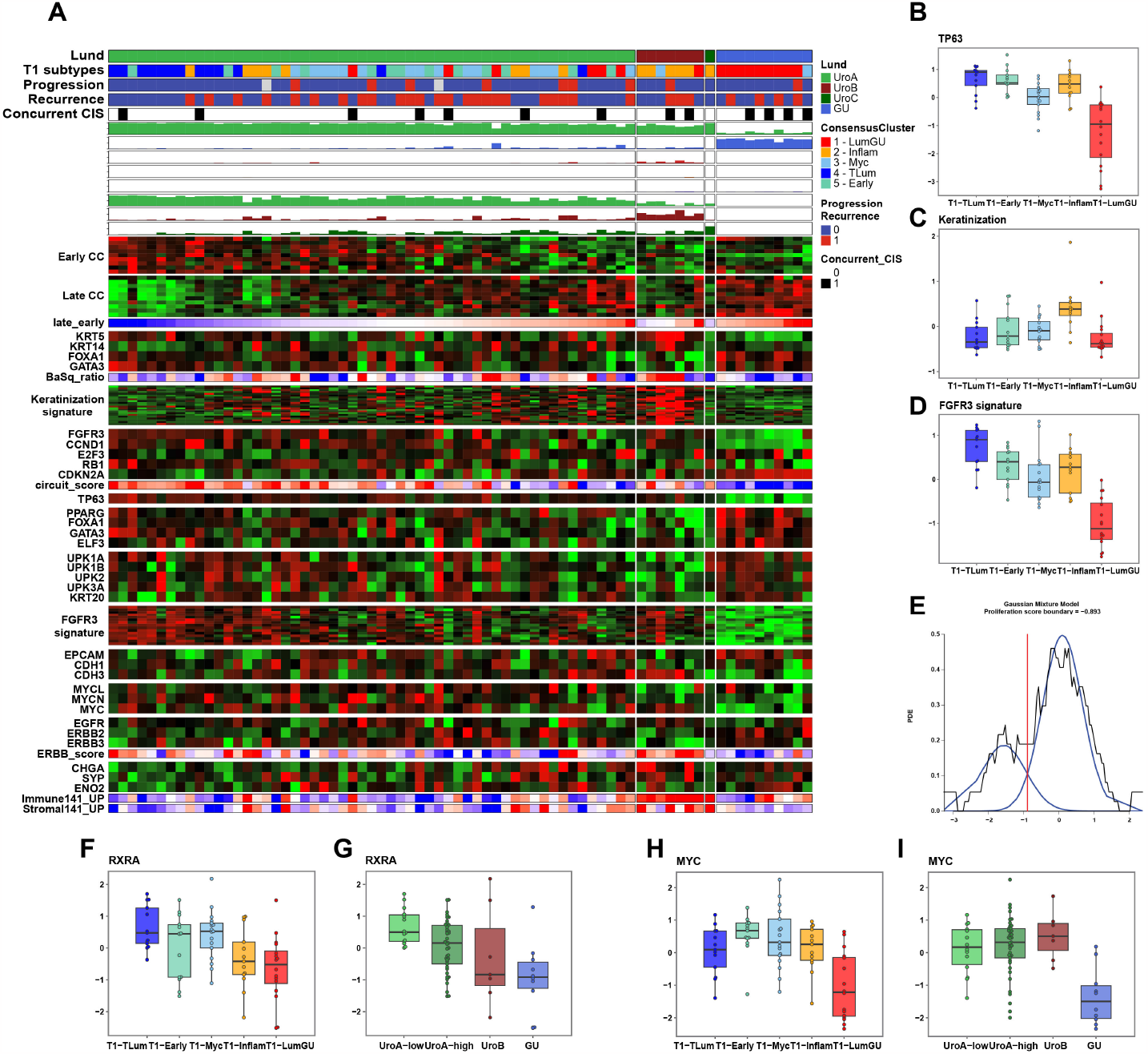
Classification of the RobertsonT1 cohort. **A:** Gene expression heatmap of the Robertson T1 cohort. Classification scores and order of gene expression signatures as in Figure 2. **B-D:** Boxplots showing expression of TP63 (**B**), the keratinization signature (**C**) and the FGFR3 signature (**D**) across the Robertson T1 subtypes. **E:**Proliferation scores of UroA samples. Black line indicates distribution of scores; blue lines, fitted distributions; red line, established threshold. **F-G:** Expression of the differentiation marker RXRA using the Robertson T1 (**F**) and the LundTax2023 (**G**). (**H-I:**) Expression of the transcription factor MYC using the Robertson T1 (**H**) and the LundTax2023 (**I**) classification.

#### BowdenT1

The Bowden cohort comprise data for 87 HG T1 cases with FFPE-extracted RNA, sequenced using 75bp paired-end reads (33). The cohort includes 23 cases with pathologically confirmed micropapillary (MP) variant histology. The authors performed unsupervised clustering and identified two major clusters of tumors, A and B, subdividing B into B1 and B2. The LundTax2023 classification of the data produced 64 Uro (74%), 21 GU cases (24%), and one Ba/Sq (1%). Subclassification of the Uro samples resulted in 51 UroA (80%), 5 UroB (8%) and 8 UroC (12.5%) (Figure 10A, Supplementary Figure 7). We noticed that UroA and UroB were assigned to the Bowden A cluster, whereas UroC and GU to the B cluster (Figure 10B). UroC is a sub-group of Uro (together with UroA and UroB), but clusters with GU when whole genome bulk RNA is used for grouping (8, 11). Even though the MP variant was enriched in the Bowden B cluster, we find it to be particularly enriched in the GU subtype that contained 15 out of the total 23 micropapillary cases. Furthermore, MP was observed in 71% of the GU cases (p *<* 10^−5^). We thus conclude that the MP variant is a distinct feature of high grade and T1 GU cases but not of the Urothelial-like subtypes UroA, UroB, and UroC.

**Fig. 10.**
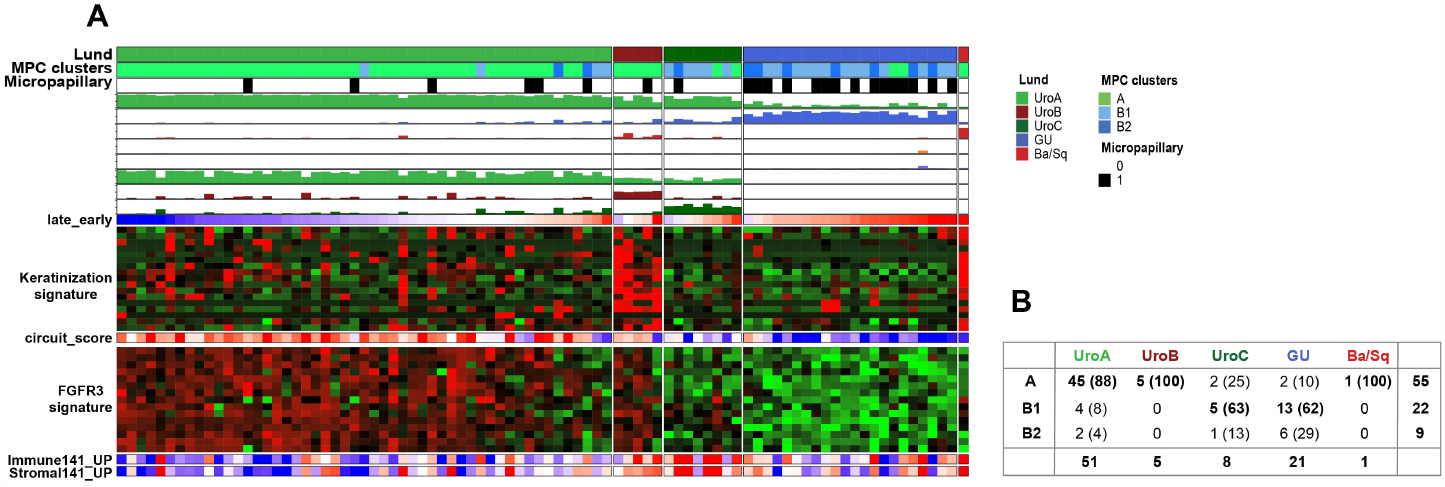
Classification of the Bowden T1 cohort. **A:** Gene expression heatmap of the Bowden T1 cohort. Classification scores and order of gene expression signatures as in Figure 2. Full heatmap is given in Supplementary Figure 7. **B:** Subtype name translation table between the Bowden T1 classification and LundTax2023 classes, n (%).

#### Summary NMIBC

The LundTax2023 almost completely recapitulated the previous classifications of the UROMOL and Leeds cohorts, showing that the updated LundTax2023 reiterates previous results (43). The analysis again indicated that the majority of cases are of the Uro subtype in these more stage-balanced NMI cohorts (Figure 11A). The analyzed T1 NMIBC cohorts were highly heterogeneous, each one selected for specific purposes, making them less comparable to each other than the MIBC cohorts analyzed previously. Still, we applied the classification algorithm and obtained results consistent with gene expression ratios and signatures used for validation. The subtype proportions in the three datasets, mainly Uro and GU cases, were similar but with a higher fraction of GU cases compared to the UROMOL and Leeds cohorts, both of which include large proportions of Ta cases (Figure 11B). It is known that GU increase in frequency in T1 tumors and predominantly are of high grade (11). Hence, selection for high grade T1 cases would select for GU cases.

**Fig. 11.**
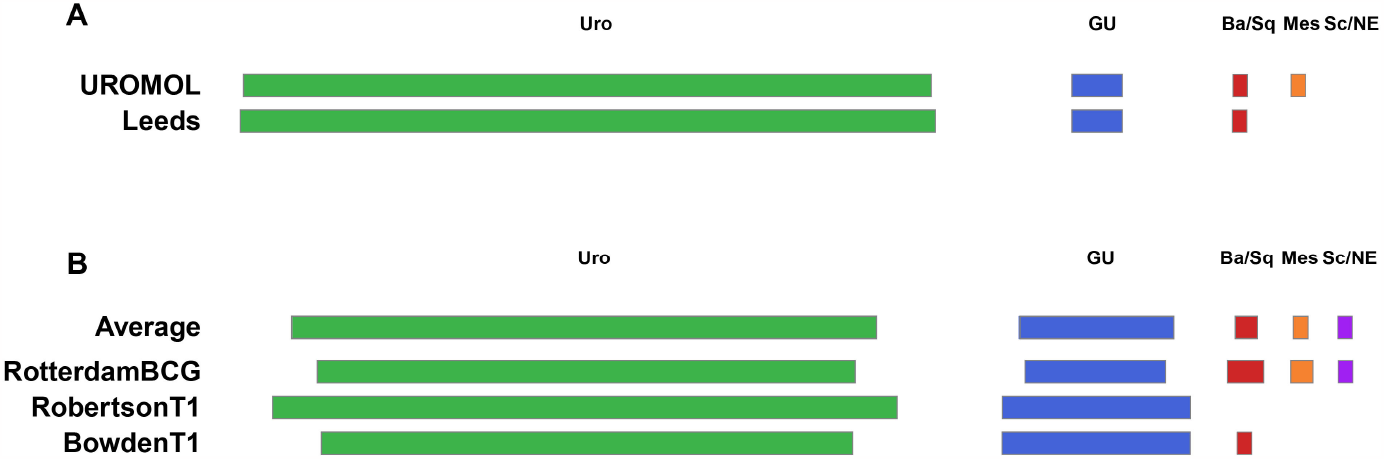
Proportion of the five-class subtypes in the individual non-muscle-invasive cohorts.

## Discussion

In the present work we developed and tested an updated cross-platform compatible rule-based prediction model for the Lund Taxonomy classification system, LundTax2023. As the source for the updating process, we used a total of four in-house datasets generated with Affymetrix microarrays or RNA-sequencing. As data preprocessing can impact the performance of a classifier, we used minimalistic preprocessing pipelines for all our training data, which can also be applied to external datasets with raw data to achieve annotation and preprocessing parity. We used RMA and fixed BrainArray annotations for the Affymetrix microarrays and Kallisto pseudoalignment with a fixed transcriptome index for the RNA-sequencing data. To further increase the robustness of the classifier we included two parallel preprocessing versions of the data when training the model. All four training datasets also had extensive IHC data in addition to RNA gene expression data. This made it possible to define the molecular subtypes in the training data according to previously determined cancer cell phenotypes, i.e., limiting subtype definitions to protein expression profiles of the cancer cells proper (8). The Lund Taxonomy differs from other systems in this respect (9). As demonstrated, the LundTax2023 RNA subtype classifier was strongly aligned with the independent IHC-based subtype classification. In addition, the classification was less affected by low tumor purity. Furthermore, the Lund Taxonomy is independent of tumor stages and is applicable as a single system for both NMIBC and MIBC including metastatic disease. The LundTax2023 classifier was validated in a large number of independent bladder cancer transcriptomic datasets, produced by different platforms, preprocessing approaches, and of different tumor stages. The subtype prediction scores were distinct across all evaluated external cohorts, with few ambiguous classification results, and recapitulated the expression patterns of the training cohort. Slightly lower and less distinct prediction scores were observed in the Seiler2019 cohort of post-chemotherapy treated RC samples, suggesting that the classifier performance may be negatively affected by either the treatment, sampling procedure, or methods of tissue preservation in this study. Despite the lower prediction confidence, the subtype assignments still captured the expected expression patterns, supporting the classification results. By this we conclude that the presented LundTax2023 algorithm may be applied to any bladder cancer transcriptomic dataset. The proportions of the five major LundTax subtypes were almost identical in the different muscle invasive cohorts indicating the stability of the algorithm. Variability was to some extent seen among the less different Urothelial-like subtypes UroA, UroB and UroC. However, the mutation data clearly showed that FGFR3 mutations were highly prevalent in UroA and UroB but almost absent in UroC. UroC was furthermore distinguished from GU by showing almost complete absence of RB1 mutations, highly prevalent in GU. We have previously described the luminal subtypes UroA, UroB, UroC and GU subtypes at the cancer cell level by extensive IHC analysis (11). The present mutation data further supports the existence of the Urothelial-like variant UroC.

A complicating factor when applying and comparing different systems is the investigators definition of subtypes and the subtype names. From a clinical point of view this may result in unreproducible, or even contradicting, results when applying different systems (46). For instance, the TCGA Basal-squamous subtype is analogous to the LundTax Ba/Sq. However, whereas the LundTax Ba/Sq is more or less strictly defined by the ratio between *KRT5, KRT14* and *GATA3* and *FOXA1* (36), the TCGA variant is more inclusive. In addition, the TCGA system does not include the well-established GU subtype (8, 11, 43, 45, 47). A comparison with the GCS system may also lead to confusing results. Even though most Uro and GU were classified as Luminal or Luminal infiltrated a large proportion were also classified as “basal”. Uro and GU are both “luminal” and there are no “infiltrated” versions in the LundTax system as infiltration is treated as a separate variable. Furthermore, the LundTax well separated Sc/NE, analogous to the TCGA Neuronal, were classified as “basal” in the majority of cases by the GSC system. However, on some occasions the translation from one system to another may be simpler as in the case of the Robertson et al. suggested system for T1 tumors, all of the suggested subtypes could be translated into LundTax equivalents. Bowden et al. performed an unsupervised clustering of high grade T1 tumors and defined A and B subtypes with no systematic comparisons with other classifications systems. The authors showed a very strong association of micropapillary histology with their subtype B. However, a reclassification of their data using LundTax2023 revealed that the A subtype corresponded to UroA and UroB and the B subtype to UroC and GU cases. The micropapillary histological variants were almost exclusively seen in the GU subtype with proportionally very few in the urothelial-like subtypes UroA, UroB and UroC, and hence making the biological background to this histological variant more precise. It has been shown before that this histological variant is associated with samples showing a GU profile (48, 49).

Two of the investigated reports describe subtype shifts as a consequence of treatment. In the Seiler2017 dataset 53% of the matched pre- and post-treatment cases changed subtype according to the GSC system, while 59% of matched cases changed subtypes after BCG treatment in the RotterdamBCG cohort according to the BRS classification system. However, when the LundTax2023 was applied the frequency of subtype changes was considerably lower, 27% and 12%, respectively. We have previously shown that recurring and progressing tumors from the same patient in most cases are of the same molecular subtype (24, 50). The reason for the discrepancy between the LundTax classification and that of Seiler et al. (1) and de Jong et al. (34) is that the LundTax system is focused on the phenotypes of the cancer cells proper and does not consider infiltration i.e., purity, as a class defining property. Most observed subtype classification changes could be attributed to altered purity of the post-treatment sample. The exercise where we introduced increasing amount of “infiltration” to the samples in silico clearly demonstrated how sensitive classifiers may be to tumor purity for stable classification. Thus, the disagreement in subtype shift frequencies is related to how molecular subtypes are originally defined and could be boiled down to the question if an infiltrated version of a subtype is a new subtype, or not (7). According to LundTax it is not. It is only by separating these variables that we can fully understand what subtype changes correspond to, and how often they occur.

With the present paper we make publicly available an updated rule-based single sample classification algorithm for the Lund Taxonomy of urothelial carcinomas. The algorithm is applicable to commonly used types of RNA expression data. As existing classification systems often differ with respect to how subtypes are defined, even if the names are similar, we believe that the most efficient approach to clinical issues is to apply and compare results obtained through different classification schemes. The LundTax2023 classifier presented here was designed exactly for these purposes.

## ACKNOWLEDGEMENTS

The authors would like to acknowledge Clinical Genomics Lund, SciLifeLab, and Center for Translational Genomics (CTG) Lund University for providing expertise and service with sequencing and analysis. We thank William Kim, Jeffrey Damrauer, Gordon Robertson, Christiaan de Jong, and Alberto Nakauma Gonzalez for their help with the respective cohorts.

## AUTHOR CONTRIBUTIONS

E.A.C. and P.E. performed bioinformatic analyses; E.A.C., P.E., G.S., and C.B. drafted the manuscript; F.L., G.S., C.B., and S.V. provided intellectual input and critically read the manuscript; F.L. provided funding. The study was conceptualized and designed by P.E.

## ETHICS STATEMENT

The study was approved by the Research Ethics Board of Lund University (2012/22; 2013/264; 2017/37).

