## Supplementary Figures for "A versatile and upgraded version of the LundTax classification algorithm applied to independent cohorts"

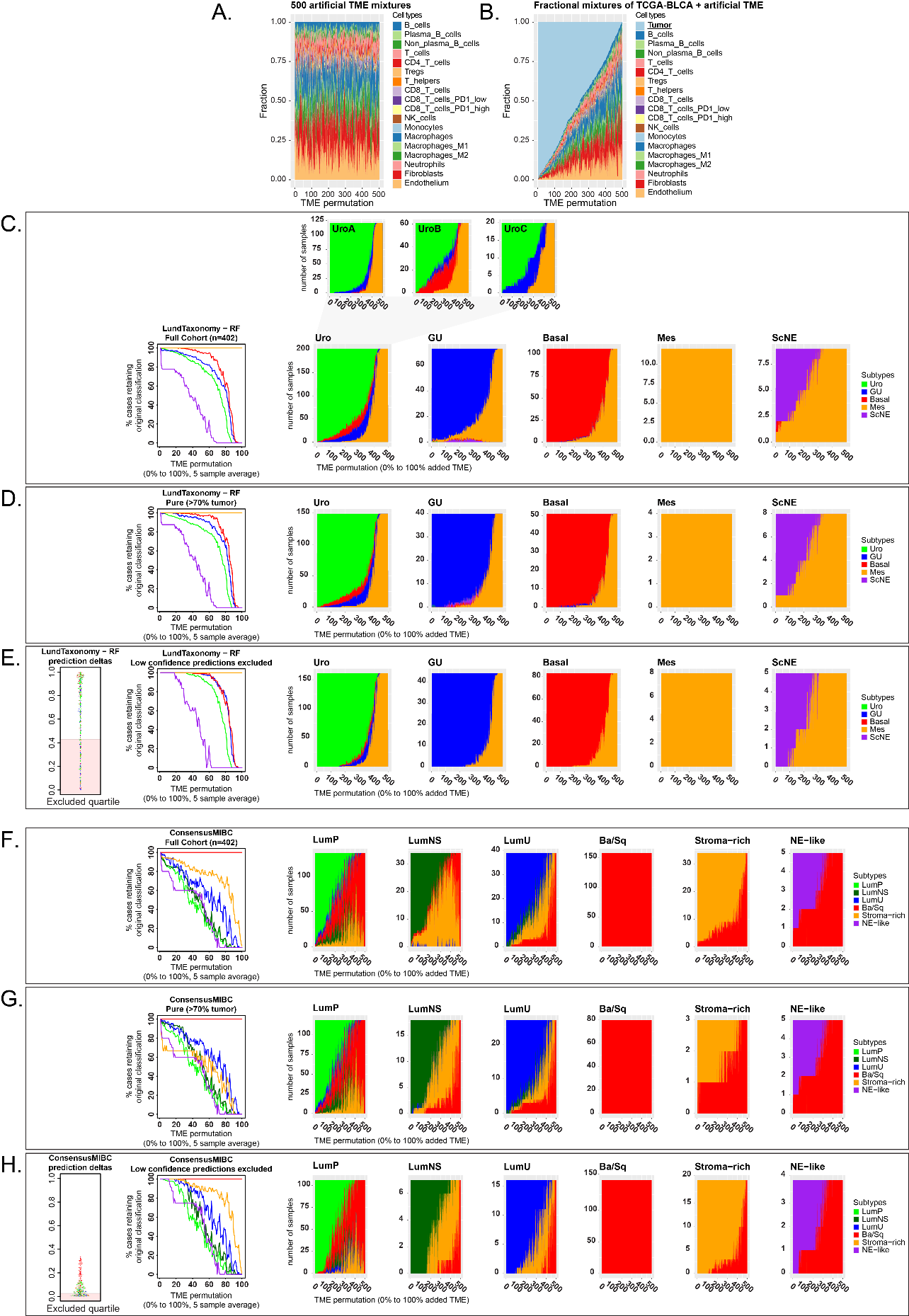

**Supplementary figure 1**. Synthetic tumor micro-environment (TME) addition to TCGA-BLCA cohort. **A:** 500 randomly generated TME cell-fractions were generated to form synthetic pure TME TPM expression profiles which were then added to 402 samples from the TCGA-BLCA dataset. **B:** The TCGA-BLCA tumor and TME composition along the 500 permutations. **C:** LundTax2023 classification results for the full TCGA-BLCA cohort (n=402) at progressively lower tumor purity. The line-plot indicates the percent of each subtype class that retains their initial classification, smoothed in a window of 5 permutation results. The stacked area graphs for each of the 5 main LundTax subtypes indicate the changing classification result across the 500 permutations of decreasing tumor purity, as illustrated in panel B. The three breakout plots for Uro indicate the 5-class switches when Uro-classified samples were stratified by UroA, UroB, and UroC subclassification. **D:** Classification results when excluding tumors already below 70% tumor purity (as assessed by the Kassandra TME deconvolution tool). The results were similar to the full cohort. **E:** Classification changes most commonly occurred in samples of lower prediction confidence at baseline. In panel E the quartile of samples with lowest prediction confidence were excluded (predicted class - second closest prediction), which reduced subtype switching. **F:** ConsensusMIBC classification results for the full TCGA-BLCA cohort (n=402) at progressively lower tumor purity. **G:** ConsensusMIBC classification results with tumors below 70% purity excluded. **H:** ConsensusMIBC classification results when excluding the quartile of samples with lowest prediction confidence (predicted class - second closest prediction).

­­
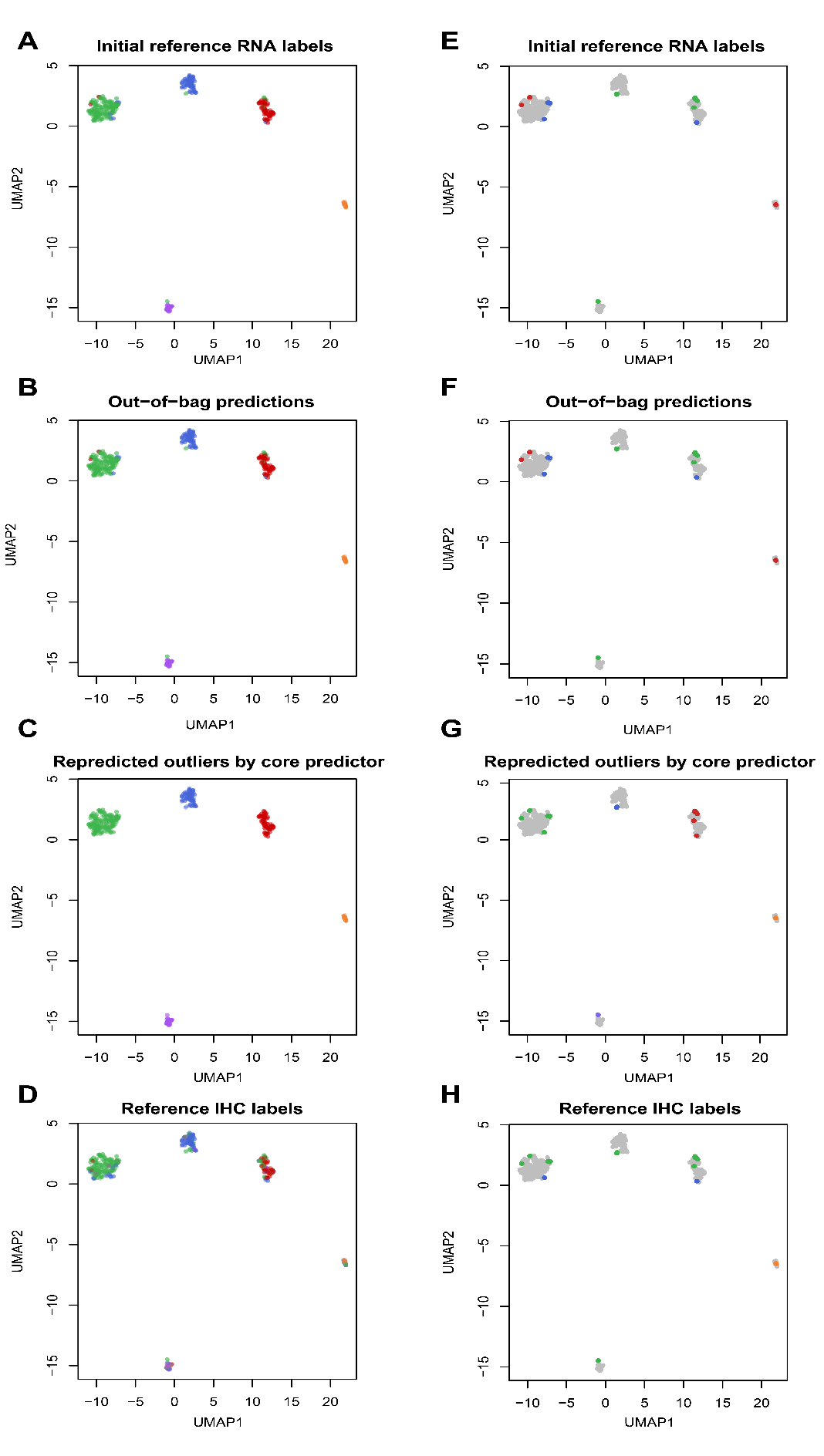

**Supplementary Figure 2.** UMAP embedding of rules used in the Lund2017 classifier. **A:** Lund2017 samples colored according to reference RNA-based labels. **B:** Lund2017 samples colored according to OOB predictions. **C:** Lund2017 samples colored according to OOB predictions, and outlier samples colored according to predictions obtained after applying a predictor trained excluding these outlier samples. **D:** Lund2017 samples colored according to IHC-based labels. **E-H:** Lund2017 samples with only outlier samples colored according to the same labels as A.B,C and D, respectively. labels as panels A,B.C and D, respectively.

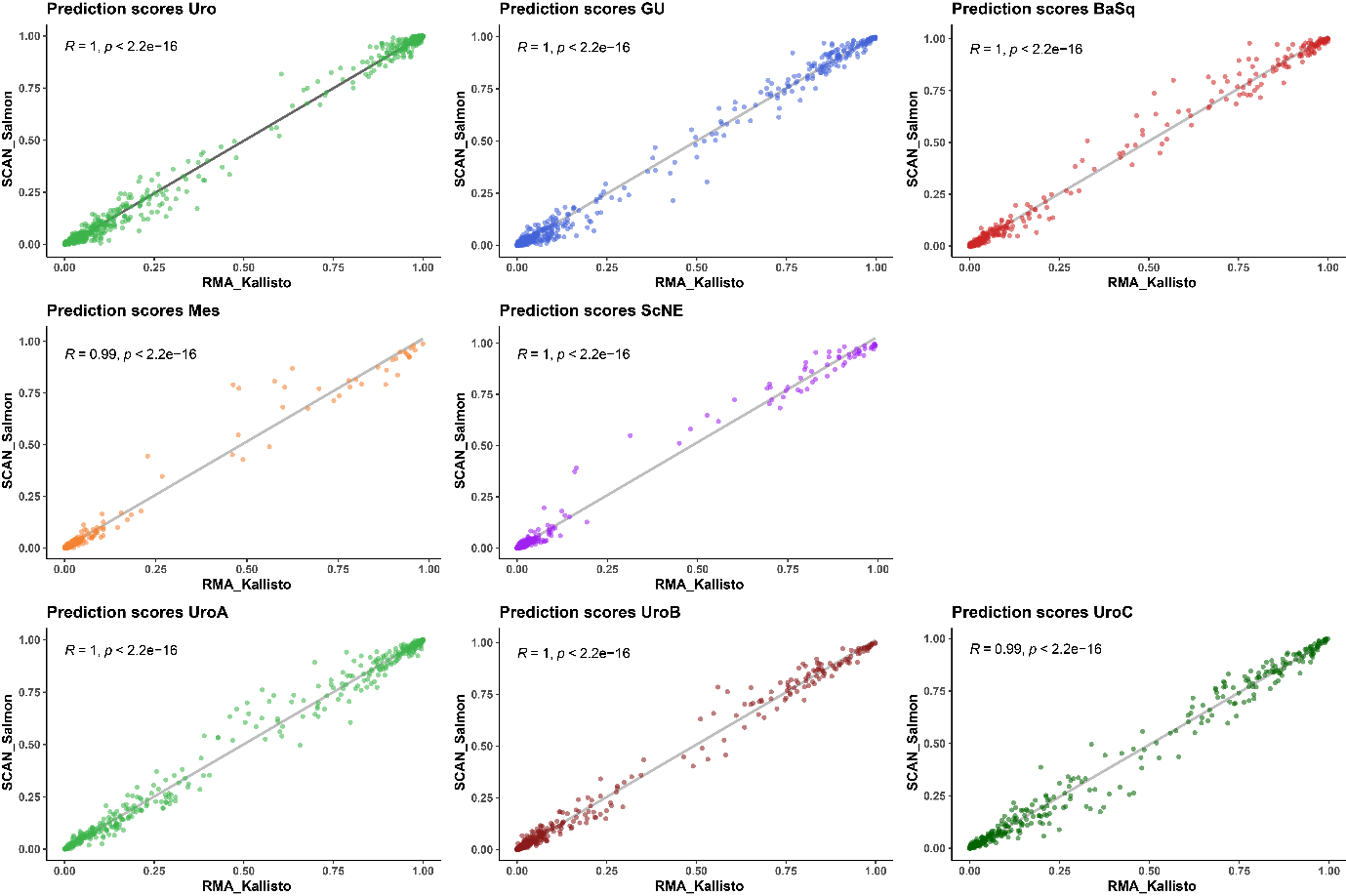

**Supplementary Figure 3.** Pearson correlation plots between different versions of the data (RMA/Kallisto vs SCAN/Salmon) for all classes. Includes all 4 datasets used in the training process (n = 1049).

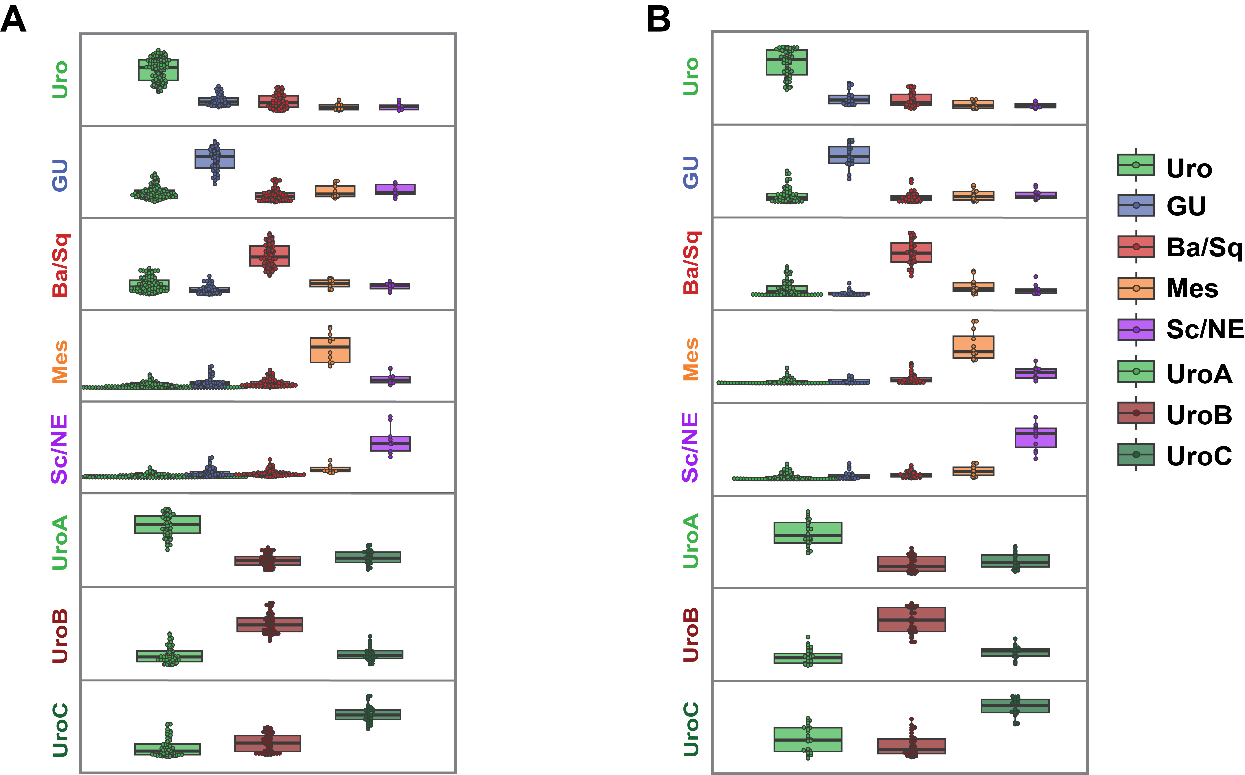

**Supplementary Figure 4.** Boxplots showing prediction scores for each class in the IMVigor210 (**A**) and UC-Genome (**B**) cohorts.

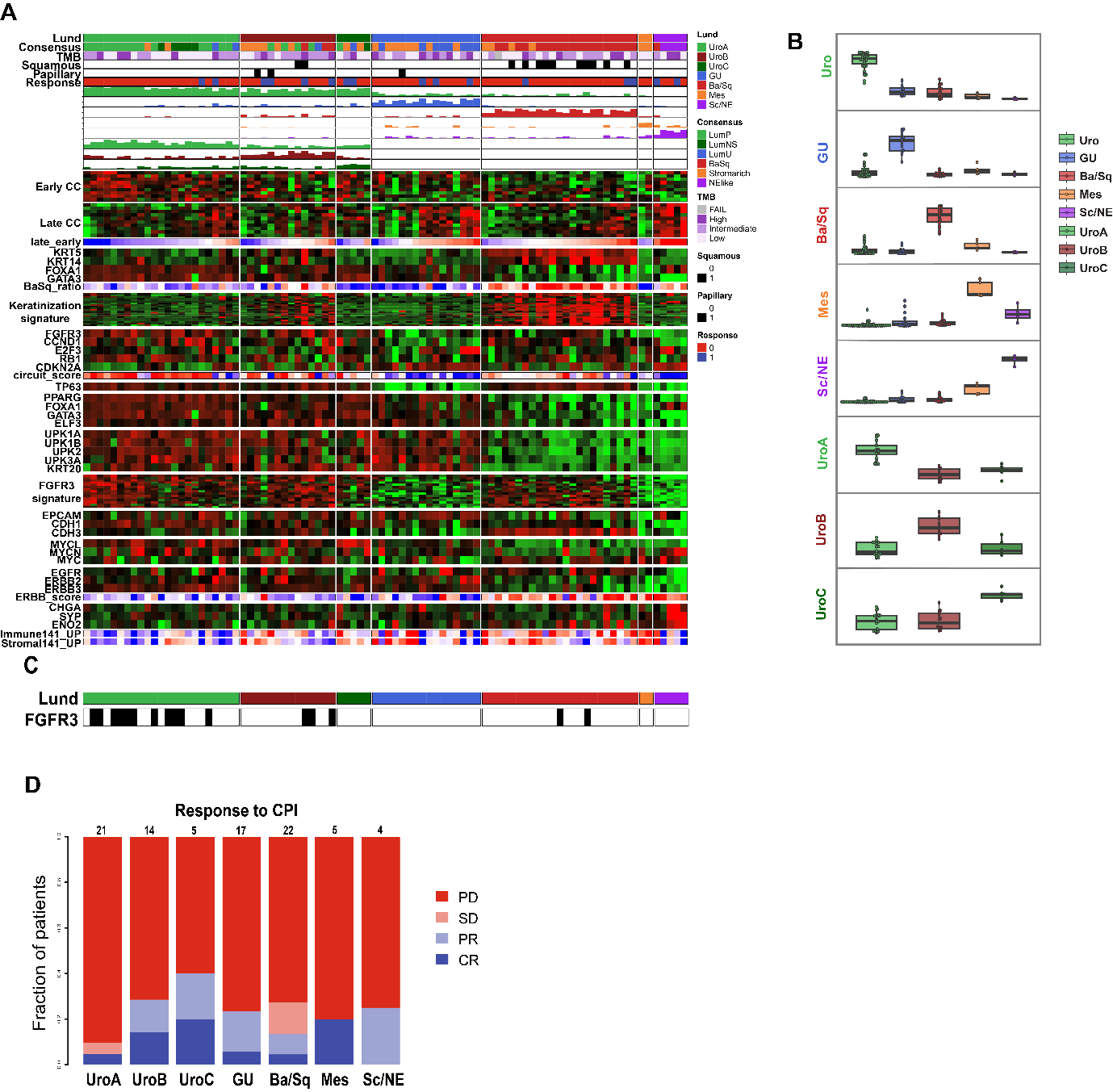

**Supplementary Figure 5.** Classification results for the UNC-108 cohort. **A:** Gene expression heatmap. Classification scores and order of gene expression signatures as in Figure 2. **B:** Boxplots of classification scores for each molecular subtype. **C:** Distribution of FGFR3 mutations. **D:** Response to checkpoint inhibitor (CPI) treatment.

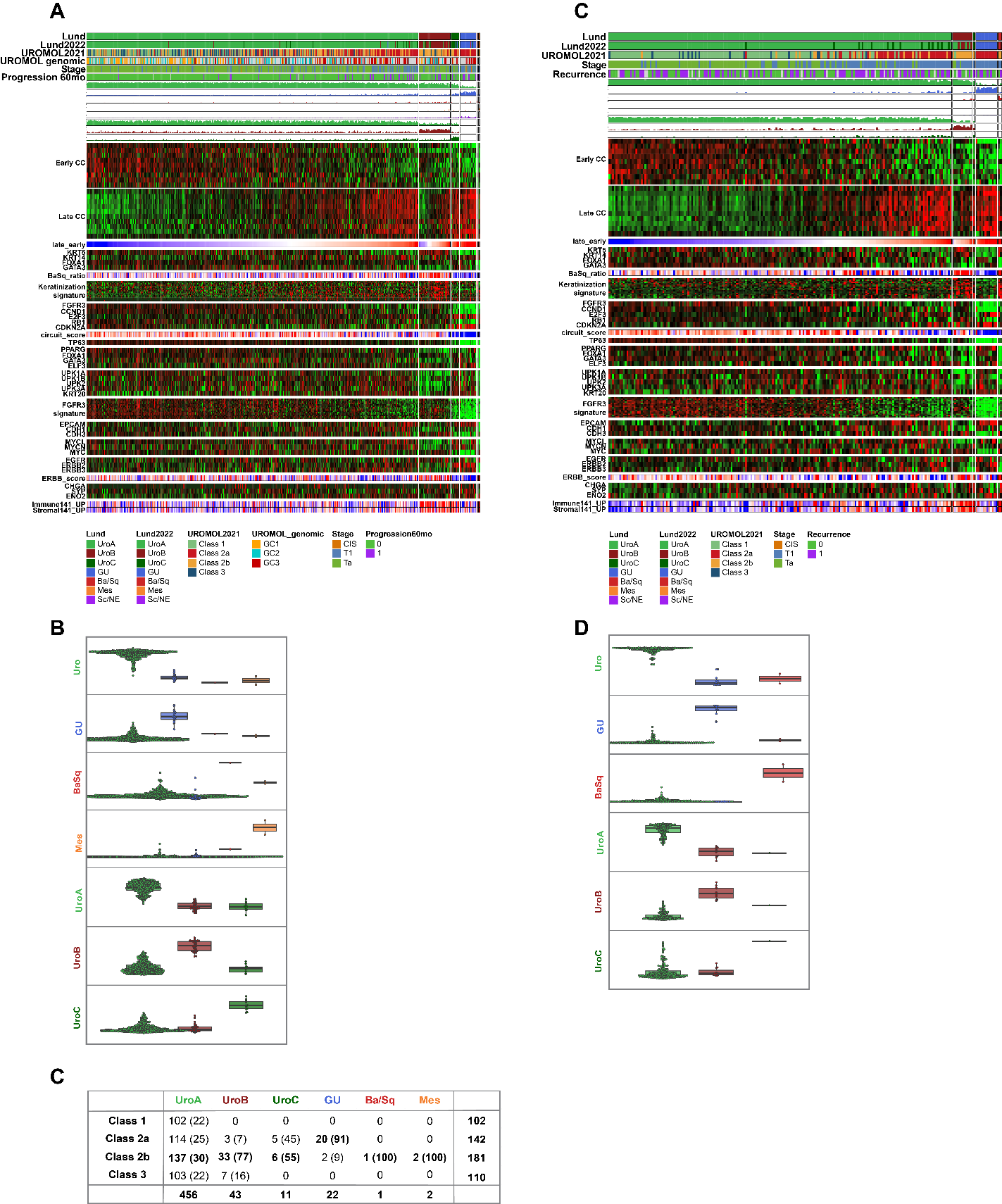

**Supplementary Figure 6.** Classification results for the UROMOL and Leeds cohorts. **A:** Gene expression heatmap of the UROMOL cohort. Classification scores and order of gene expression signatures as in Figure 2. **B:** Boxplots of classification scores for each molecular subtype in the UROMOL cohort. **C:** A subtype name translation table between the UROMOL and LundTax2023 classes. **D:** Gene expression heatmap of the Leeds cohort. Classification scores and order of gene expression signatures as in Figure 3. **E:** Boxplots of classification scores for each molecular subtype in the Leeds cohort.

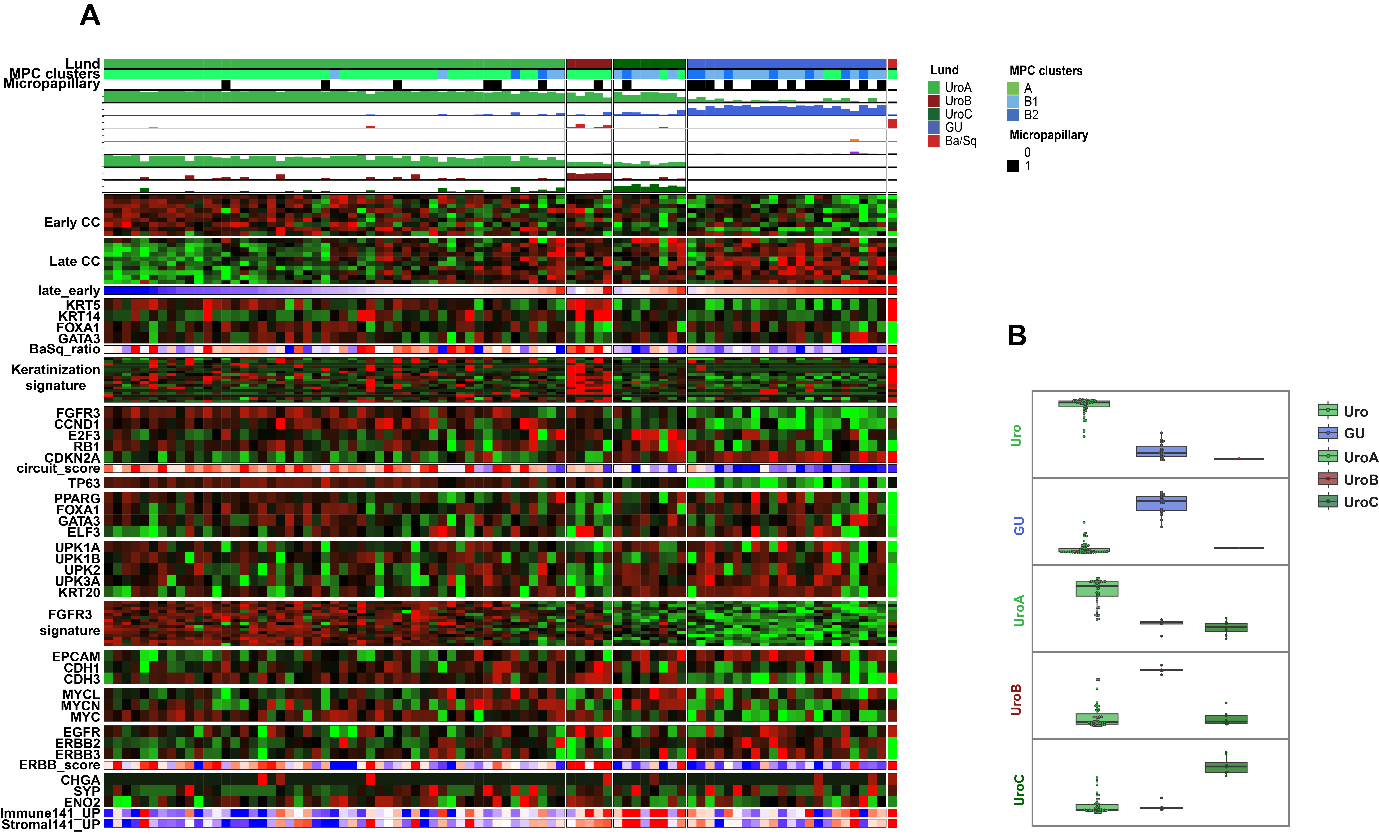

**Supplementary Figure 7.** Classification results for the Bowden T1 cohort. **A:** Gene expression heatmap. Classification scores and order of gene expression signatures as in Figure 2. **B:** Boxplots of classification scores for each molecular subtype.
