## Supplementary Table 2 for "A versatile and upgraded version of the LundTax classification algorithm applied to independent cohorts"

| Cohort | Platform | Number of samples | Database | Database ID | Reference |
| --- | --- | --- | --- | --- | --- |
| **Lund2017** | Affymetrix Human Gene 1.0 ST Array | 307 | GEO | GSE83586 | *1* |
| **Lund2020** | Affymetrix Human Gene 1.0 ST Array | 173 | GEO | GSE128959 | *2* |
| **Lund2022** | Affymetrix Human Gene 1.0 ST Array | 310 | GEO | GSE169455 | *3* |
| **Lund265** | Illumina HiSeq | 265 | Zenodo | 10362517 | 4 |
| **TCGA-BLCA** | Illumina HiSeq | 402 | GDC | TCGA-BLCA | *5* |
| **TCGABiolinks** | Illumina HiSeq | 431 | TCGABiolinks R package (GDC) | TCGA-BLCA | *5* |
| **TCGA Toil** | Illumina HiSeq | 426 | USCS Toil Xena Hub | TCGA-BLCA | *5* |
| **TCGA recount3** | Illumina HiSeq | 433 | Recount3 R package | TCGA-BLCA | *5* |
| **Seiler2017** | Affymetrix Human Exon 1.0 ST Array | 305 | GEO | GSE87304 | *6* |
| **IMVigor210** | Illumina HiSeq | 347 | EGA | EGAS00001004343 | *7* |
| **Seiler2019** | Affymetrix Human Exon 1.0 ST Array | 133 | GEO | GSE124305 | *8* |
| **Bowden T1** | Illumina HiSeq | 87 | GEO | GSE136401 | *9* |
| **Robertson T1** | Illumina HiSeq | 73 | GEO | GSE154261 | *10* |
| **UNC-108** | Illumina HiSeq | 89 | GEO | GSE176307 | *11* |
| **Leeds** | Affymetrix Human Transcriptome Array 2.0 | 217 | GEO | GSE163209 | *12* |
| **UROMOL** | Illumina HiSeq | 535 | EGA | EGAS00001004693 | *13* |
| **UC-Genome** | Illumina HiSeq | 176 | dbGaP | phs003066.v1.p1 | *14* |
| **RotterdamBCG** | Illumina HiSeq | 179 | BRSpred R package  and EGA (raw data) | EGAS00001006879 | *15* |

**Supplementary Table 2.** Description of collected datasets. Name of the cohort as referred to in the text, gene expression platform, number of samples, database where the data is located, dataset ID in the correspondent database, and reference publication, are indicated for all the published datasets. For the TCGA data, 4 versions of the dataset were collected through different portals.

1 Sjödahl G, Eriksson P, Liedberg F, et al. Molecular classification of urothelial carcinoma: global mRNA classification versus tumour-cell phenotype classification. J Pathol. 2017;242:113–125.

2 Sjödahl G, Eriksson P, Patschan O, et al. Molecular changes during progression from nonmuscle invasive to advanced urothelial carcinoma. Int J Cancer. 2020;146:2636–2647.

3 Sjödahl G, Abrahamsson J, Holmsten K, et al. Different Responses to Neoadjuvant Chemotherapy in Urothelial Carcinoma Molecular Subtypes. European Urology. 2022;81:523–532.

4 P. Eriksson, E. Aramendia, G. Sjödahl, et al. Lund Bladder Cancer Group - Lund Taxonomy 2023 Classifier - Training Data. Zenodo, 2023.

5 Robertson AG, Kim J, Al-Ahmadie H, et al. Comprehensive Molecular Characterization of Muscle-Invasive Bladder Cancer. Cell. 2017;171:540-556.e25.

6 Seiler R, Ashab HAD, Erho N, et al. Impact of Molecular Subtypes in Muscle-invasive Bladder Cancer on Predicting Response and Survival after Neoadjuvant Chemotherapy. Eur Urol. 2017;72:544–554.

7 Mariathasan S, Turley SJ, Nickles D, et al. TGFβ attenuates tumour response to PD-L1 blockade by contributing to exclusion of T cells. Nature. 2018;554:544–548.

8 Seiler R, Gibb EA, Wang NQ, et al. Divergent Biological Response to Neoadjuvant Chemotherapy in Muscle-invasive Bladder Cancer. Clin Cancer Res. 2019;25:5082–5093.

9 Bowden M, Nadal R, Zhou CW, et al. Transcriptomic analysis of micropapillary high grade T1 urothelial bladder cancer. Sci Rep. 2020;10:20135.

10 Robertson AG, Groeneveld CS, Jordan B, et al. Identification of Differential Tumor Subtypes of T1 Bladder Cancer. Eur Urol. 2020;78:533–537.

11 Rose TL, Weir WH, Mayhew GM, et al. Fibroblast growth factor receptor 3 alterations and response to immune checkpoint inhibition in metastatic urothelial cancer: a real world experience. Br J Cancer. 2021;125:1251–1260.

12 Hurst CD, Cheng G, Platt FM, et al. Stage-stratified molecular profiling of non-muscle-invasive bladder cancer enhances biological, clinical, and therapeutic insight. *Cell Rep Med*. 2021;2(12):100472

13 Lindskrog SV, Prip F, Lamy P, et al. An integrated multi-omics analysis identifies prognostic molecular subtypes of non-muscle-invasive bladder cancer. Nat Commun. 2021;12:2301.

14 Damrauer JS, Beckabir W, Klomp J, et al. Collaborative study from the Bladder Cancer Advocacy Network for the genomic analysis of metastatic urothelial cancer. Nat Commun. 2022;13:6658.

15 de Jong FC, Laajala TD, Hoedemaeker RF, et al. Non-muscle-invasive bladder cancer molecular subtypes predict differential response to intravesical Bacillus Calmette-Guérin. Sci Transl Med. 2023;15:eabn4118.
